# Uncovering novel therapeutics for schizophrenia: a multitarget approach using the CANDO platform

**DOI:** 10.1101/2025.10.24.684257

**Authors:** Yakun Hu, Katherine Elefteriou, Zackary Falls, Ram Samudrala

## Abstract

Schizophrenia is a complex and debilitating neuropsychiatric disorder characterized by positive, negative, and cognitive symptoms, many remaining insufficiently addressed by current treatments. High rates of treatment resistance and heterogeneous pathophysiology pose significant challenges to traditional single-target drug discovery. To address this, we applied the CANDO platform to identify repurposable drugs for schizophrenia using a multitarget strategy. The platform evaluates how compounds interact with the entire human proteome, generating interaction signatures that capture a compound’s effect across all targets. By comparing these signatures, CANDO computes compound similarity scores and enables consensus prediction of novel therapeutics. Across all indications and for schizophrenia in particular, CANDO outperformed controls by orders of magnitude across several benchmarking metrics, accurately recovering known drug–indication relationships. We applied our CANDO pipeline, benchmarked by its ability to recover approved drugs for their indications using similarity- and consensus-based metrics, to generate a ranked list of repurposable compounds. Our comprehensive literature review confirmed clinical or biochemical evidence supporting 25 high-corroboration drug candidates, including phenothiazine and benzamide antipsychotics, tricyclic antidepressants, benzodiazepines, and monoamine oxidase inhibitors. We identified protein targets with the highest likelihood of interaction with these top drugs and assessed prediction quality through overlap analysis with gold standards, showing significantly better concordance with established schizophrenia-related biology than controls from bottom-ranked and randomly selected drugs. Among these proteins are corroborated targets such as canonical neurotransmitter receptors of the dopamine, serotonin, and adrenergic classes, as well as monoamine transporters, tyrosine aminotransferase, and lysine-specific histone demethylase 1A. Enrichment of calcium-binding proteins among the top predicted targets for the phenothiazine antipsychotic thiethylperazine highlights a potential role for dysregulated calcium signaling, including calmodulin–CaMKK2 pathways, in schizophrenia pathology and treatment response. Another phenothiazine antipsychotic, triflupromazine, binds the serotonin transporter in addition to its canonical dopamine D2 receptor interaction, highlighting its potential relevance to depressive symptom modulation in schizophrenia. These findings demonstrate the utility and accuracy of the CANDO platform in elucidating multitarget pharmacological mechanisms and accelerating the identification of effective repurposable treatments for schizophrenia.

## Introduction

Schizophrenia is a complex and debilitating psychiatric disorder characterized by psychotic symptoms such as persistent hallucinations and delusions, negative symptoms including diminished emotional expression, avolition, and social withdrawal, and cognitive deficits such as poor attention, memory, and executive function^1–5^. Affected individuals die on average 28.5 years earlier due to comorbid medical conditions and suicide^2,6^. Its average lifetime prevalence is approximately 1% of the population.^1^. Antipsychotic medications are the cornerstone of schizophrenia treatment, alleviating positive symptoms such as hallucinations and delusions by reducing dopaminergic hyperactivity through dopamine D2 receptor blockade in mesolimbic pathways. Treatment resistance remains common, even though second-generation antipsychotics are generally more effective than first-generation antipsychotics at alleviating cognitive and negative symptoms through enhanced serotonergic 5-HT2 receptor antagonism and modulation of dopaminergic activity in mesocortical pathways^1,5,7–9^. An estimated one-third of patients have treatment-resistant schizophrenia. As a result, they experience little or no improvement despite adequate trials of multiple antipsychotics^5^. Additionally, long-term remission of symptoms is achieved in only approximately one third of patients^6^. Adverse effects of antipsychotics include hyperprolactinemia and an elevated risk of diabetes mellitus and cardiovascular-related mortality. First-generation antipsychotics carry a high risk of extrapyramidal side effects such as Parkinsonian tremors and tardive dyskinesia, while second-generation antipsychotics can increase weight gain^3^. According to Reimers et al., the anticholinergic and antihistaminic effects of some antipsychotics account for deficits in processing speed, verbal memory, and executive functions^10^. Overall, novel therapeutic strategies are needed to address the broad symptom profile of schizophrenia, overcome treatment resistance, and minimize side effects.

Traditional drug discovery is a costly and time consuming endeavor due in part to the need to perform separate biochemical analyses of potentially millions of compounds^11^. As a result, computational strategies, benefiting from advances in computing power and the expanding availability of molecular data, including ligand properties, binding interactions, and binding-site dynamics informed by target 3D structures, have become indispensable for accelerating the identification of new drug candidates^11,12^. A key strategy is virtual high-throughput screening that uses computer simulations to evaluate large libraries of compounds against a biological target and significantly reduces cost by computationally triaging compounds prior to pharmacological testing^11^. Two main protocols used in virtual screening are structure-based virtual screening and ligand-based virtual screening. In structure-based virtual screening, computational methods dock candidate small molecules into the target’s binding site to predict their binding strength and orientation. Ligand-based virtual screening leverages known active ligands for a target along with the principle that molecules with similar structures or properties are likely to exhibit similar biological activities^13^. Recent efforts to improve upon conventional virtual screening protocols include hybrid and multistage screening pipelines where structure- and ligand-based virtual screening can be merged in sequential or parallel workflows to leverage their respective strengths^14^. Recent advances in AI co-folding models, such as AlphaFold 3, which can directly predict protein–ligand structural complexes, provide improved binding-site conformations that can enhance structure-based and hybrid virtual screening workflows^15^.

The Computational Analysis of Novel Drug Opportunities (CANDO) is a platform that enables multitarget and multiscale drug repurposing and discovery by ranking compounds for specific indications based on their interaction profiles across the proteome^16,17^. CANDO is based on the principle of polypharmacology, screening drugs across the full ensemble of potential on and off targets simultaneously, rather than focusing on a single target as in traditional structure-based virtual screening^16,18^. The central hypothesis is that drugs with similar compound–proteome interaction signatures will exhibit similar biological or therapeutic effects^16,19^. This enables the discovery of functionally similar drugs that may be missed by ligand-based virtual screening. The predictive pipelines in CANDO have been experimentally validated across a range of indications, demonstrating its ability to propose clinically relevant drug–disease connections^17,18,20–24^. In this study, we apply the CANDO platform to identify new potential drug candidates for treating schizophrenia, predict the most likely protein targets of these drugs to elucidate their potential mechanisms of action, and validate these predictions against curated gold standard protein sets.

## Methods

### Overview of schizophrenia drug repurposing pipeline using the CANDO platform

We initiated our repurposing pipeline by curating compound, protein, and indication libraries to form the basis for compound–proteome interaction scoring. Interaction scores between each compound and protein were generated using an in-house scoring protocol, which uses predicted binding sites and ligand–compound fingerprint similarity to produce proteomic interaction signatures. CANDO compares these signatures across all compounds to create ranked similarity lists. We then used these lists in benchmarking to evaluate performance in recapturing known drug–indication associations. We ranked compounds by their frequency of appearance and rank across similarity lists of known drugs to predict potential schizophrenia treatments. We manually curated the top 100 predictions through a literature search to identify 25 high-corroboration repurposing candidates based on clinical and biochemical evidence. Control sets were generated by extracting structurally complex compounds from the bottom of similarity rankings and by selecting random entries. Next, we identified predicted protein targets for each compound in all prediction and control sets. To evaluate prediction quality, we curated gold standards from PubMed / DrugBank, UniProt, and GeneCards against which target lists were compared using overlap analyses.

### Drug and protein library construction

The CANDO pipeline begins by curating libraries of drugs, proteins, and indications^25^. We used a DrugBank-sourced compound list with a filter to include only the 2,449 drugs that are approved by at least one regulatory agency, such as the FDA or its international counterparts. We used the Homo sapiens AlphaFold2 library as the proteome, consisting of 20,295 AlphaFold2 structure-predicted proteins^25^. We used the Comparative Toxicogenomics Database to link 2,449 approved drugs to 22,771 drug-indication associations, identifying drugs by DrugBank ID and indications by Medline Subject Headings terms^25–27^. Interaction scores between drugs and proteins were computed across the full protein library to generate a drug–proteome interaction matrix. We used this matrix, along with the drug-indication mappings, for benchmarking performance and predicting new therapeutic candidates.

### Drug-proteome signature generation

We calculated drug-drug distances based on proteomic interaction signatures for each small molecule drug in the CANDO library and used them to benchmark the platform and generate predictions. Our in-house bioanalytical docking protocol (BANDOCK) computes interaction scores between each drug and every protein in the human proteome by integrating predicted binding site information from COACH and chemical similarity^19^, with higher scores predicting greater interaction probability, with a maximum of 1.0^20,23,25,28,29^. Chemical similarity was computed by comparing the COACH-predicted ligand to the query compound using RDKit-generated Extended Connectivity Fingerprints with a diameter of 4, with similarity quantified by the Sorenson-Dice coefficient^20,25^. BANDOCK calculates a vector of interaction scores known as the interaction signature^25^ for each drug to generate the drug-proteome interaction matrix. Cosine distances were calculated between drug-proteome interaction signatures in an all-against-all fashion to assess drug similarity. This resulted in ranked similarity lists for each drug that were used to benchmark known drug-indication associations and to predict novel treatment options^25^.

### Benchmarking

We benchmarked CANDO for schizophrenia and across all indications using four metrics against random and hypergeometric controls to validate its predictive accuracy and ranking quality. Indication accuracy (IA) measures how often the platform correctly identifies drugs approved for the same indication as similar using a leave-one-out approach, calculating the percentage of drug similarity lists that include at least one other approved drug within a pre-determined cutoff (e.g., top 10 or top 25)^23,30,31^. The platform calculates average indication accuracy (AIA) by computing the IA for every indication mapped to at least two drugs, the minimum for determining recovery in a similarity list, and averaging the results^31^. New indication accuracy (nIA) measures how well the platform recaptures associated drugs in the consensus list of an indication. It is defined as the percentage of drugs withheld from their indication that appear in the consensus list for that indication within a given cutoff. New average indication accuracy (nAIA) is the mean nIA across every indication^25^. CANDO uses the ranking metric normalized discounted cumulative gain (NDCG), commonly applied in information retrieval to evaluate the quality of ranked lists based on known relevant items, to assess how well the platform prioritizes drugs for specific indications and across all indications^25^. The discounted cumulative gain (DCG) metric assigns higher weight to correctly indicated drugs appearing near the top of the ranked similarity lists, reflecting the principle that early-ranked relevant items contribute more to the overall relevance score^30–32^. CANDO computes ideal DCG (IDCG) from a hypothetical perfect ranking where all approved drugs are placed at the top of the similarity list, serving as the reference for computing the normalized score^25,30^. NDCG is calculated by dividing a DCG by the IDCG and yields normalized values between 0 and 1, with higher scores indicating better ranking^30^. The platform uses a related metric, new NDCG (nNDCG), to measure prediction quality and is calculated from consensus rankings rather than similarity lists. We compared our results for each benchmarking metric to controls generated by taking the average of 100 randomized drug–protein interaction matrices to model the platform’s expected performance under random chance.

### Generating putative drug predictions

We used the CANDO canpredict module to generate a list of putative drug candidates to treat schizophrenia. The module predicts candidates by counting how often a compound appears in the top 10 of similarity lists for drugs already approved for the indication. It prioritizes drugs that appear more frequently and resolves ties by favoring those with better average ranks across the lists. For each drug, probability was calculated as the likelihood that a drug would appear among the top 10 ranked compounds in similarity lists as many times as observed under random ranking, computed using a binomial cumulative density function. We selected the top 100 drugs from this ranking for subsequent literature review. As noted earlier, we limited our predictions to approved drugs to ensure acceptable safety profiles.

### Literature search to corroborate putative drug candidates

We conducted a literature search using PubMed and Google Scholar to corroborate our drug predictions, using queries that combined “schizophrenic” or “schizophrenia” and each drug name, limited to the title or abstract fields. We marked as “high-corroboration” drugs with one high-corroboration and one low-corroboration evidence or two high-corroboration evidences. A high-corroboration evidence is defined as 1) a clinical study that demonstrates the efficacy of the drug as a standalone treatment for schizophrenia 2) a functional assay that demonstrates agonism or antagonism at a relevant receptor, or 3) a binding study that shows the drug has potent binding activity to a relevant receptor. A low-corroboration evidence is defined as a clinical study that demonstrates the combined efficacy of the drug as an adjunctive treatment along with another drug. We marked as “refuted” drugs that were once approved but have since been taken off the market due to lack of efficacy. All other drugs were marked as “no data found”.

### Generating bottom-ranked and random controls

From the similarity list of each schizophrenia-associated drug, we exclude structurally simplistic molecules that have fewer than 4 heavy (non-hydrogen) atoms. The drugs with the most appearances in these bottom lists were selected as the bottom filtered predictions control. We generated the random control by selecting random ranks from the similarity list of risperidone, a top-selling antipsychotic for treating schizophrenia, and extracting the drugs from the corresponding rows.

### Top protein interactions for putative drug candidates

We applied CANDO’s target ranking module to generate a list of 100 top protein predictions for each of our high-corroboration drug predictions. This module ranks proteins for a given drug based on their interaction scores, returning the targets with the strongest predicted associations. We repeated this process to generate control target lists for the bottom filtered and random drug lists.

### Curation of gold standard libraries

We curated the UniProt gold standard library by obtaining protein functional annotations for each schizophrenia associated drug present in UniProt and compiling the unique results. UniProt returns entries with evidence-supported binding annotations derived from its own manual curation, linking each drug to proteins whose ligand-binding sites have been experimentally validated. Because these annotations are standardized through Chemical Entities of Biological Interest identifiers and supported by literature or structural data, the resulting protein set reflects biologically grounded drug–target interactions. Next, we curated the GeneCards gold standard library by applying three layers of filters. First, we restricted results to the sections “Disorders, Function, Genomics, Pathways, Phenotypes, Proteins, and Publications.” Following application of this filter, the dataset was enriched for biologically relevant associations and curated to remove entries derived largely from automated text mining or other low-evidence sources^33^. Lastly, we removed genes that do not code for proteins or those with relevance scores of less than ten, to cull the list from around 8,000 to just under 70. A combined gold standard library between PubMed and DrugBank was curated by using the UniProt recommended names of the top targets of our high-corroboration predictions and the names of every drug for which that target was a top prediction in CANDO. We included in the gold standard drug-protein pairs with evidence of a direct binding interaction between them, excluding indirect interactions such as downstream effectors. Following the PubMed review, we conducted a DrugBank search for each high-corroboration drug to identify proteins involved in its mechanism of action. We curated the library by including proteins that overlap with those found in the CANDO top target list for that drug. The results from the PubMed and DrugBank literature search are presented in Figure 4, Table 2, and the discussion section of our paper as well as Supplementary Table 2.

### Functional target overlap analyses between high-corroboration drug target lists and gold standards

We compared each of the top, bottom, and random predicted target lists against each gold standard to obtain metrics of similarity; results are shown in figure 3. We calculated relative frequencies as the number of intersections between a target list and a gold standard within each of five rank bins, divided by the total number of intersections with that gold standard. These histograms highlight which rank ranges contribute most to the overlap and help assess prediction quality. We calculated overlap percentage as the cumulative number of intersecting proteins up to a given rank cutoff, divided by the total number of proteins in the gold standard list. We used this metric to compare the list of top targets of high-corroboration drug predictions against controls. The Jaccard coefficient, defined as the size of the intersection divided by the size of the union between the gold standards and prediction sets, was used to quantify and compare their overall similarity.^34,35^ The higher the Jaccard coefficient, the greater the similarity, and the better our prediction list captures proteins independently corroborated by gold standard libraries.

### Inclusion and exclusion criteria for Discussion

From our 25 high-corroboration drug predictions, we excluded from our analyses 12 that were formerly FDA approved and marked as discontinued, but not due to efficacy or side effect reasons^36^, or are approved for schizophrenia treatment in Europe, Canada, and other international jurisdictions. Since the efficacy of these drugs is already known, we did not highlight them as prominent repurposing candidates. An exception is made for drugs that have recent evidence of activity at a protein other than the ones associated with their originally marketed effects. Of the remaining, we identified drugs that were highly ranked, within the top 50, of our consensus predictions. Among these, we included drugs that had literature-backed interactions with protein targets, as documented in Table 2.

### Exploratory analysis on sources of interaction scores for high-ranking targets of phenothiazines

We observed that canonical targets of thiethylperazine, our fourth ranked prediction, were absent from its top 100 CANDO targets. We investigated its mechanism of action by running the CANDO top targets module to extract the highest likelihood protein interactions for thiethylperazine and seven schizophrenia associated drugs that included it within their similarity lists, then formed a consensus ranking by ordering proteins by their frequency of appearance across these lists. We repeated this procedure for three comparison cohorts: seven non-phenothiazine antipsychotics from our predictions, eleven non-antipsychotic schizophrenia predictions, and fifteen random approved DrugBank drugs, and compared functional characteristics across sets. For each predicted drug–protein interaction, we extracted the top-scoring Protein Data Bank (PDB) ligand associated with that interaction from BANDOCK. A literature review was conducted through PDB for each ligand-target pair to determine the mechanism of action of predicted interactions and to corroborate our assessment of therapeutic relevance. Predicted targets of phenothiazines that could be traced to a top ligand that was also a phenothiazine and with literature evidence of being implicated in schizophrenia are highlighted in Table 3 and were added to our PubMed/DrugBank gold standard set (see our code repo on Dryad).

## Results and Discussion

### Benchmarking results

Figure 2 presents benchmarking results for schizophrenia (Figure 2 A/B) and across all indications (Figure 2 C/D), showing recapture percentages based on similarity and consensus lists as well as prioritization of indicated drugs. In panel A, schizophrenia indication accuracy (IA) results show recapture rates ranging from 75% at the top 10 cutoff to 93% at the top 100 cutoff, while new IA (nIA) increases from 10% to 53%, both markedly better than controls. This indicates the ability of CANDO to recover known schizophrenia associated drugs based on interaction signature similarity and thereby reflects its predictive strength. Panel B reports normalized discounted cumulative gain (NDCG) values for schizophrenia ranging from 0.151 at the top 50 cutoff to 0.220 at the top 10, while new NDCG (nNDCG) ranges from 0.058 at the top 10 to 0.138 at the top 100. NDCG and nNDCG values outperformed the average matrix control across all benchmarking cutoffs, highlighting CANDO’s ability to prioritize indicated drugs for schizophrenia compared to random chance. Panel C highlights all indications IA (AIA) performance between 22% and 44%, a greater than five-fold improvement on control values at the top 10 cutoff. New AIA (nAIA) ranges from 9% to 26%, compared to control values between 0.7% and 6.8%. These findings indicate that CANDO consistently identifies and ranks drugs approved for the same indication as highly similar to one another across all indications. In panel D, NDCG values across all indications span 0.044 to 0.059, while controls remain between 0.005 and 0.014; average nNDCG for all indications ranges from 0.049 to 0.083, outperforming control values of 0.003 to 0.014 across all cutoffs. These results highlight the accuracy of the CANDO platform in recapturing and ranking putative drug candidates for schizophrenia.

**Figure 1.**
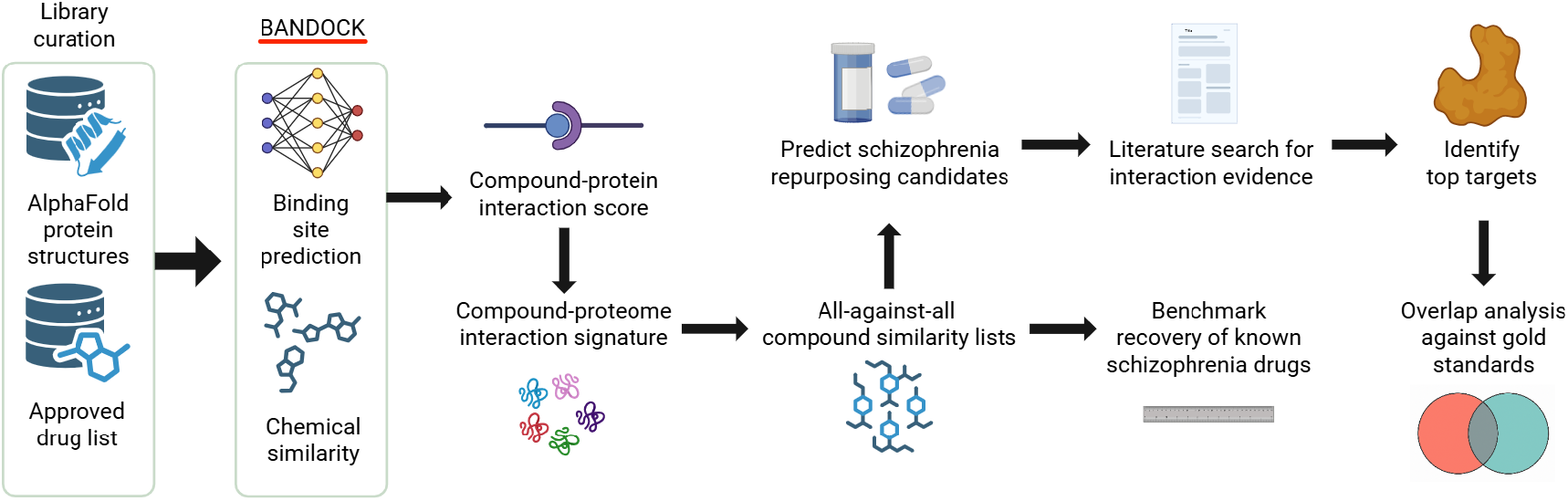
Overview of the CANDO pipeline for generating and corroborating drug repurposing candidates for schizophrenia. We used a protein library of AlphaFold-predicted structures and an approved list of drugs from DrugBank to generate drug-protein interaction scores using our in house bioanalytical docking (BANDOCK) protocol. Interaction scores were computed across the full proteome to produce an interaction signature for each drug. Drug-drug similarity is computed using cosine distance between interaction signatures in an all-against-all comparison. We used these similarity rankings to benchmark the platform and to predict repurposing candidates. Top-ranked drug predictions were corroborated through literature review, and their most likely predicted protein targets were compared to gold standard protein sets via functional target overlap analysis. This workflow summarizes the CANDO platform’s computational pipeline for generating drug–proteome interaction signatures, benchmarking drug–drug similarity, and identifying repurposable schizophrenia therapeutics through functional target validation.

**Figure 2.**
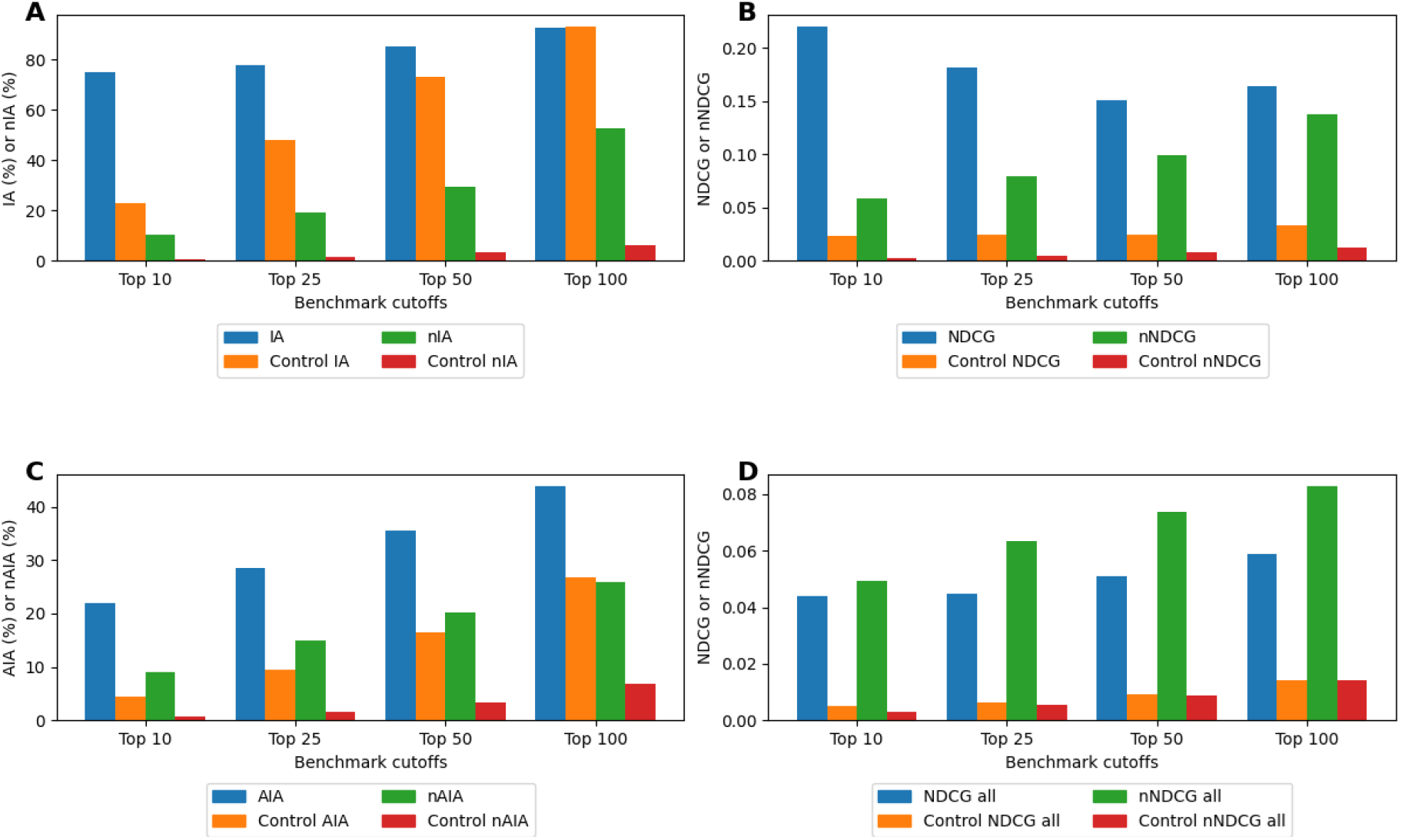
Benchmarking performance of the CANDO platform for schizophrenia and across all indications. We report benchmarking metrics that measure how effectively known drug–indication associations are recovered and ranked for schizophrenia and for all indications. (A) Schizophrenia IA (blue) and nIA (green) are plotted across top-k cutoffs (Top 10, 25, 50, 100) and compared with their respective random-control values (orange and red). IA for schizophrenia outperforms the control by over three-fold at the Top 10 cutoff and remains better until the Top 100, where both exceed 90%, while nIA maintains more than an eight-fold improvement across all cutoffs. (B) Schizophrenia NDCG (blue) and nNDCG (green) are displayed at the same cutoffs and compared to their corresponding control metrics (orange and red). NDCG is nearly five times better than the control at the Top 100 cutoff, with relative advantage increasing at stricter cutoffs, and nNDCG exceeds its control by more than ten-fold at the Top 100 and nearly thirty-fold at the Top 10. (C) AIA (blue) and nAIA (green) across all indications are shown across top-k cutoffs alongside their random-control values (orange and red). AIA reaches 44% at the Top 100 cutoff and significantly exceeds the control, while nAIA reaches 26%, representing an approximately four-fold improvement over the control. (D) Mean NDCG (blue) and nNDCG (green) across all indications are plotted across cutoffs with their control counterparts (orange and red). Both metrics outperform the random control by roughly four-fold (NDCG) and six-fold (nNDCG) at the Top 100 cutoff, with increasingly strong advantages at more stringent cutoffs. These benchmarking results demonstrate CANDO’s ability to recover known drug–indication relationships and prioritize relevant compounds near the top of its consensus rankings.

### Putative drug candidates and drug-indication literature corroboration

The efficacy of each putative drug candidate was evaluated through a comprehensive literature review using PubMed and Google Scholar. Table 1 lists the candidates with high-corroboration identified in the literature. Among the 100 putative drug candidates, 25 demonstrated strong clinical evidence for efficacy against schizophrenia, yielding a 25% hit rate. Of these 25, 12 are known treatments for schizophrenia, discontinued in the U.S. or approved internationally, lending credibility that the platform is able to predict structurally related and established drugs. The list includes drugs from multiple chemical and mechanistic classes, including phenothiazine antipsychotics, benzodiazepines, tricylic antidepressants, benzamide antipsychotics, and N-arylpiperazines. CANDO’s polypharmacological insight across these drug classes is supported by Table 2, which shows multitarget actions of several high corroboration drug predictions. The top 100 putative drug candidate predictions for schizophrenia can be found in Supplementary Table S1.

**Table 1.**
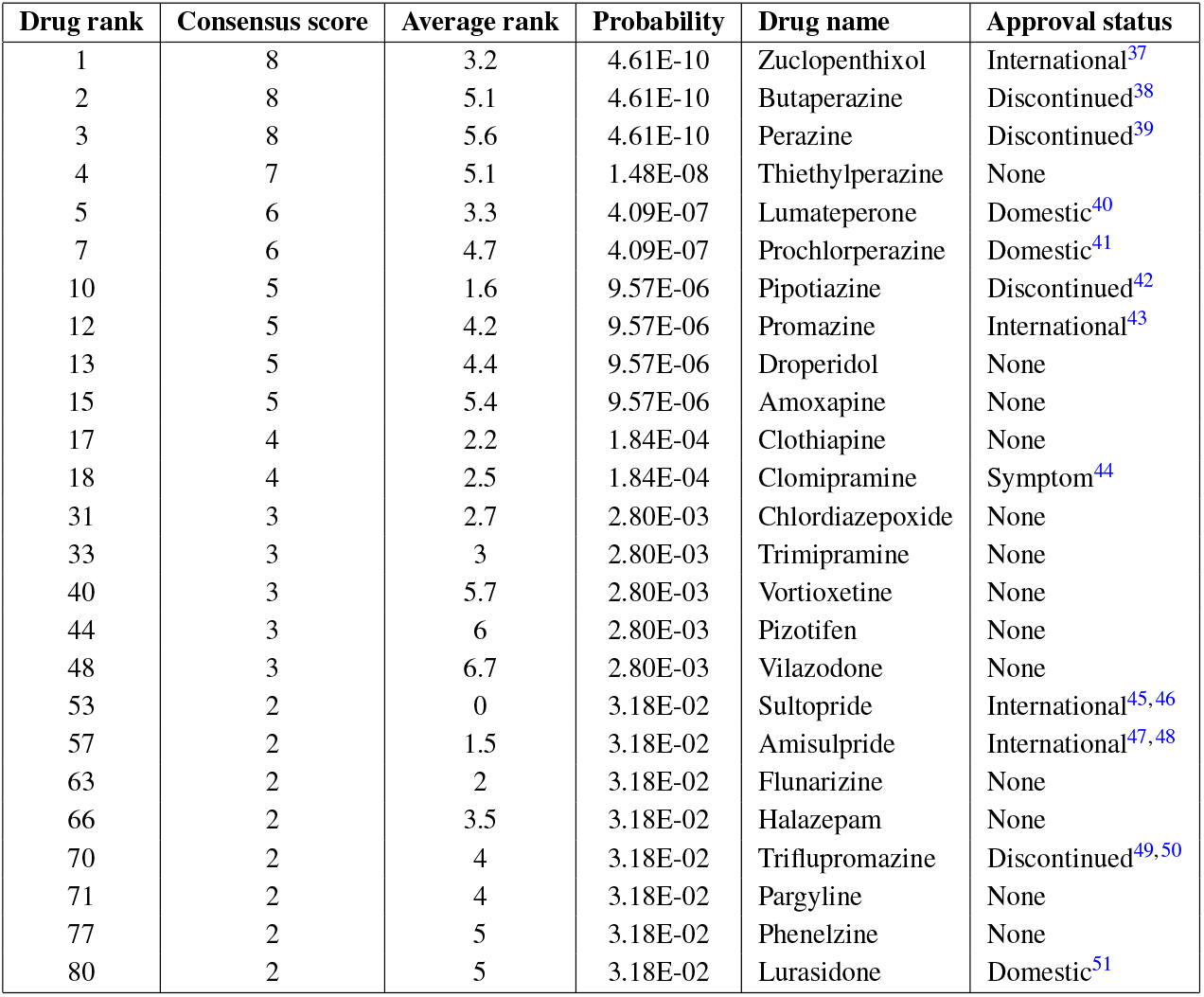
High-corroboration putative drug candidate predictions for schizophrenia derived from CANDO consensus ranking and validated through literature review. CANDO ranks drugs in descending order of consensus score, the number of times they appeared within a specified cutoff (here, n = 10) in the similarity lists of drugs associated with schizophrenia. Ties are broken by increasing order of average rank of the candidate in these similarity lists. CANDO not only predicts known antipsychotic therapies for schizophrenia but also uncovers less obvious candidates including pargyline, a monoamine oxidase inhibitor, and pizotifen, an antamine and migraine medication. Clomipramine is approved for the treatment of a schizophrenia symptom. These results indicate that the CANDO platform successfully identifies drug candidates for schizophrenia that are corroborated by existing literature and reinforce its predictive validity for drug repurposing.

**Table 2.**
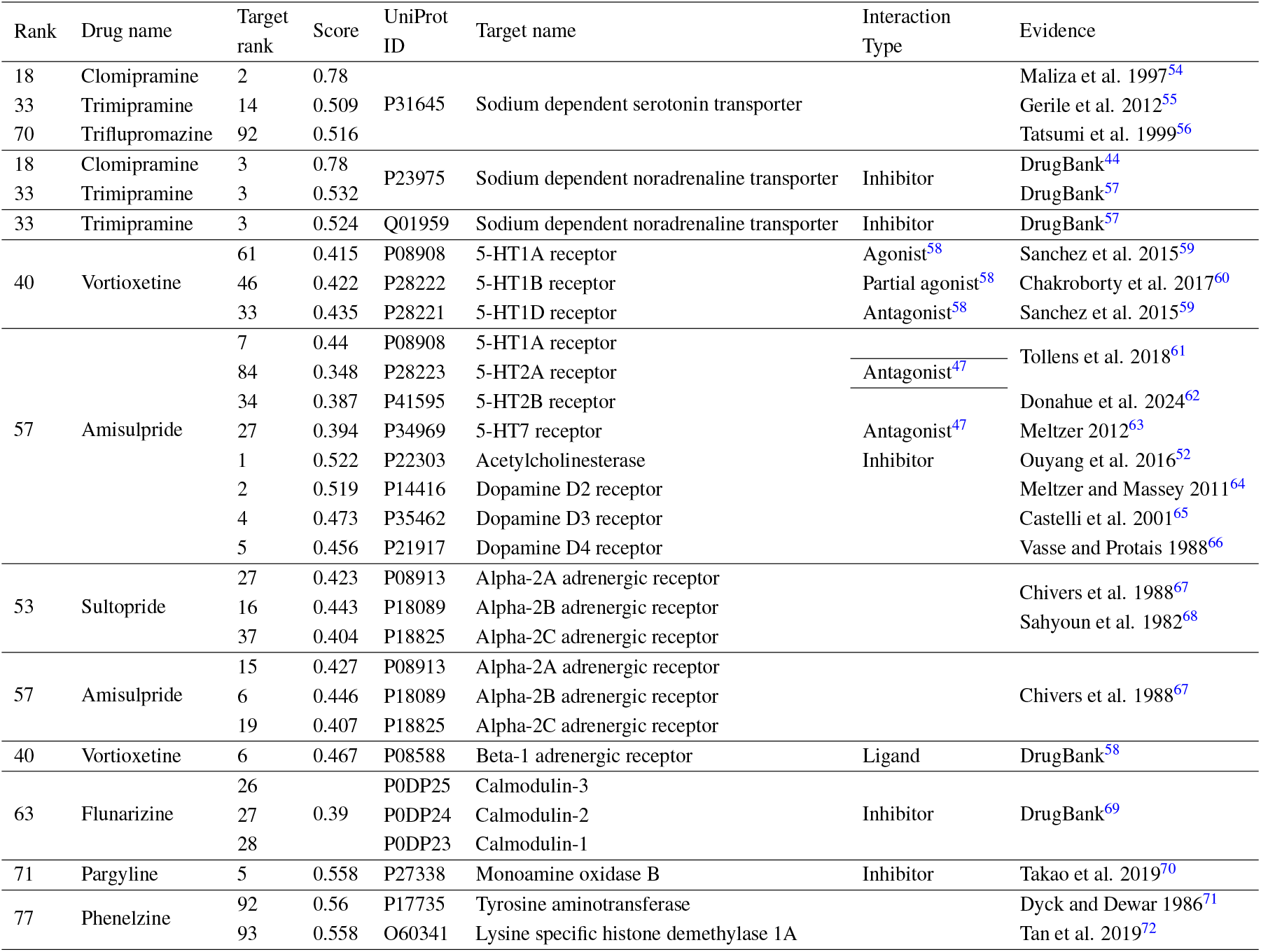
Literature corroborated top protein targets of putative drug candidates for schizophrenia. Drug rank refers to the position of the drug in our consensus-based predictions. Target rank is the position of the protein in the top targets list of the drug. Score represents the computed BANDOCK interaction score between the drug and protein. Beyond the well-established serotonin and dopamine transporters and receptors, we highlight monoamine transporters, adrenergic receptors, calmodulin, monoamine oxidase B, and acetylcholinesterase as being worthy of further investigation. See Supplementary Table S2 for the full set of target predictions for our top 25 drugs.

**Table 3.**
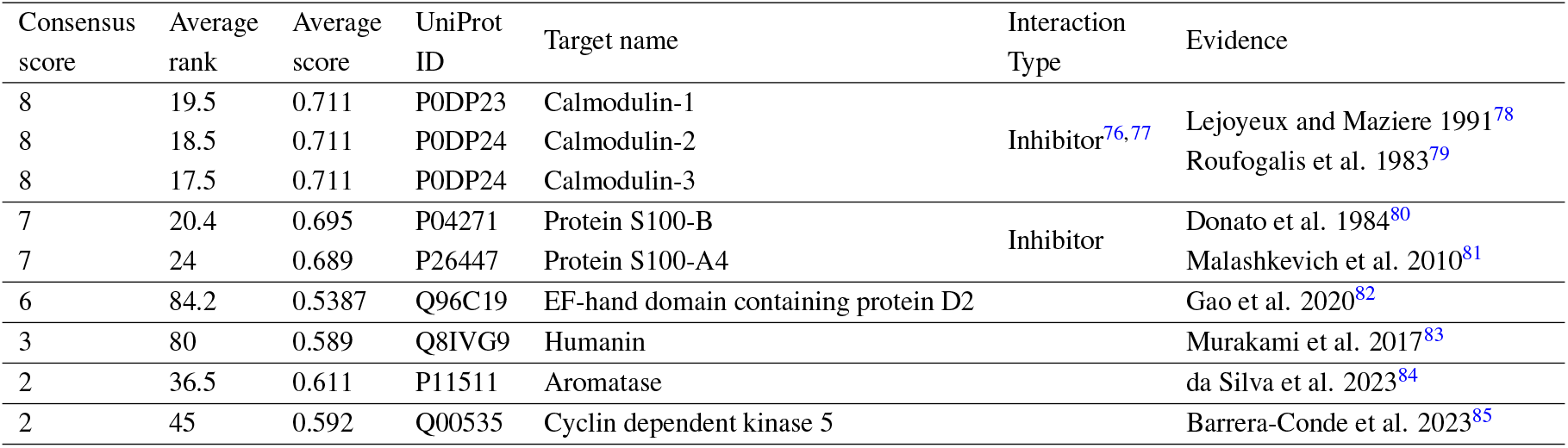
Literature corroborated top protein targets of thiethylperazine and seven schizophrenia associated drugs that include it in their similarity lists. All eight drugs, namely thiethylperazine, acetophenazine, mesoridazine, trifluoperazine, periciazine, thioridazine, perphenazine, fluphenazine, are phenothiazine antipsychotics. Proteins are ranked by the number of appearances across the eight lists; average rank is the mean position of a protein on the lists where it appears, and average score is the mean predicted interaction probability across those same lists. We retained proteins that appear on a consensus top targets list across the eight drugs and have literature evidence of involvement in schizophrenia such as differential expression. All but the bottom three proteins are calcium binding. Notably, calmodulin mediates many calcium-dependent processes in the brain, and its dysregulation could contribute to connectivity deficits observed in schizophrenia, supporting its candidacy as a therapeutic target^73^. S100B has likewise been proposed as a treatment target, as elevated concentrations may be toxic, whereas deletion or inhibition yields beneficial effects^74,75^. This convergence on calcium-binding proteins provides novel support for investigating them as central mediators of both therapeutic effects and adverse outcomes of phenothiazine antipsychotics.

#### Literature-supported protein targets identified from top CANDO drug–protein predictions for schizophrenia

We identified 29 protein targets from the top target lists of the top 25 drug predictions that were corroborated by the literature. These include canonical neurotransmitter receptors (serotonin, dopamine, adrenergic), monoamine transporters, and calcium-binding proteins. Several drugs exhibited notable polypharmacology: amisulpride, with receptor interactions extending beyond its primary targets, and vortioxetine, spanning serotonergic and adrenergic systems. We also observed secondary predictions, or targets not typically emphasized in the primary drug profile. Examples include amisulpride’s acetylcholinesterase inhibition that may confer cognitive effects^52^, and phenelzine’s histone demethylase activity, suggesting an epigenetic role beyond well-documented monoamine oxidase inhibition^53^. These findings highlight CANDO’s capacity to capture the multitarget activity central to effective schizophrenia treatments, while secondary predictions can motivate new hypotheses for exploring polypharmacology in both schizophrenia and depression.

### Overlap analyses

In Figure 3A, approximately 47% of overlaps between the top targets from high-corroboration drug predictions and the PubMed+DrugBank gold standard, 28% of overlaps with UniProt, and 33% of overlaps with GeneCards fall within the highest-ranked bin (ranks 1–20) (Figure 3). The outperformance of the random control in the highest-ranked bin is readily explained. Seven of the thirteen targets in this bin interact with the phenothiazine antipsychotic prochlorperazine or the serotonin norepinephrine reuptake inhibitor venlafaxine. If both of these drugs were excluded from our random set, the height of the leftmost random control bar would fall below that of the targets from top drug predictions bar. The GeneCards gold standard includes the dopamine and serotonin transporters. Venlafaxine targets both, skewing the overlap with the random set that has only six overlaps in total. Similarly, with an absolute frequency of only four overlaps in the bottom filtered control, the apparent enrichment in the highest-ranked bin is of limited interpretative value. Figure 3B: The overlap percentage of targets from top drug predictions ranges from approximately 55% to 100% in the PubMed and DrugBank gold standard, 6% to 19% for UniProt, and 7.5% to 18% for GeneCards. The consistent increase in overlap with rank cutoff indicates that experimentally supported targets, those overlapping with the gold standards, are distributed across the ranked list, supporting the use of larger cutoffs for validation of predicted targets. Figure 3C: Whereas overlap percentage increased with rank cutoff, reflecting the accumulation of additional experimentally supported targets, the Jaccard coefficient declined for PubMed and GeneCards as lower-ranked predictions were added. This trend indicates that while more targets are recovered at larger cutoffs, the relative enrichment declines because the total number of unique targets (the denominator of the Jaccard coefficient) increases faster than the number of overlaps, suggesting that higher-ranked predictions are of greater quality. Meanwhile, the unchanged Jaccard coefficient values observed at all rank cutoffs for the targets from the random and bottom control groups in the PubMed/DrugBank and GeneCards gold standards confirm that the observed enrichment is not due to chance.

**Figure 3.**
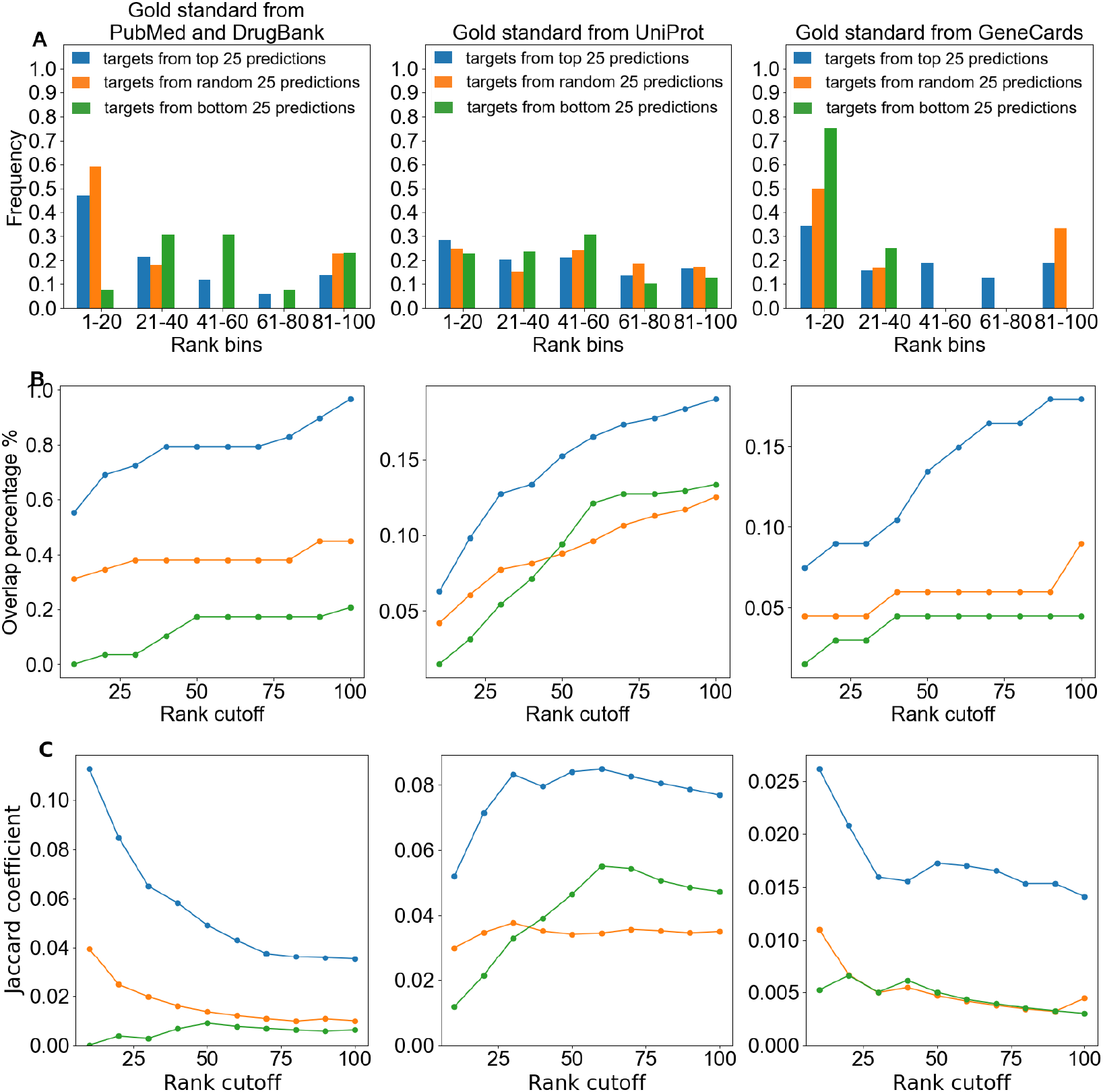
Enrichment analyses of top targets from high-corroboration drug candidates versus random and bottom-ranked controls against literature and database-derived gold standards. In all 9 charts, the blue lines or bars represent overlaps between the predicted targets from top 25 predictions and each gold standard library, green represents overlaps with targets from bottom 25 predictions, and orange represents overlaps with targets from random 25 predictions. (A) For all three gold standards, overlaps with top targets from high-corroboration drug predictions are most enriched in the top rank bin (1–20), indicating that overlaps are not randomly distributed and that higher-ranked predictions are more likely to be meaningful. (B) A greater percentage of targets from our top 25 drug predictions overlap with gold standards than the bottom and random controls across all rank cutoffs. The top 100 overlap percentage for the targets from top predictions is nearly 20% for both UniProt and GeneCards, while that of the controls stayed below 12.5% and about 9%, respectively. These results indicate that on the whole, our top target predictions for high-corroboration drugs are more enriched for biologically meaningful targets, supporting the validity of the drug rankings generated by CANDO. (C) The Jaccard coefficient demonstrates better performance across all rank values for top targets from high-corroboration drug predictions compared to controls across all three gold standards, reinforcing the biological relevance and consistency of CANDO’s target rankings. The signal for targets from top 25 predictions is concentrated at the top ranks for PubMed/DrugBank and GeneCards, whereas progression into the lower ranks introduces increasing noise. In contrast to random or bottom-ranked sets, the clear enrichment of top targets from top drug predictions confirms that CANDO’s rankings prioritize biologically relevant associations.

Our ranked predictions revealed strong recovery of known antipsychotics as well as several promising candidates with biological relevance to schizophrenia. In the following paragraphs, we discuss selected compounds that met the inclusion and exclusion thresholds outlined in the Methods.

Several of our predictions address symptoms of the disorder associated with serotonergic dysregulation, including depression and cognitive symptoms, as depicted in Figure 4. The phenothiazine antipsychotic **triflupromazine** demonstrated binding activity at the serotonin transporter (SERT)^50,86^ (Fig 4). Clinical studies show that triflupromazine significantly improves thinking disorganization, psychiatric ward behavior, and total morbidity scores on the Psychotic Reaction Profile compared with a sedative control^87^. Its potency as a SERT binder suggests potential effects on depressive symptoms, but further research is needed to confirm this, particularly in the context of schizophrenia. Triflupromazine was discontinued by the FDA, but not due to efficacy or side effect reasons^49^.

The tricyclic antidepressant (TCA) **trimipramine** also demonstrated inhibitory activity at SERT^55^. Accordingly, Eikmeier et al. provided evidence that the drug significantly decreased the anxiety / depression factor of the Brief Psychiatric Rating Scale^88^, making it a strong candidate for addressing serotonin dysregulation in schizophrenia, often linked with negative symptoms of the disorder. Chronic administration of a selective serotonin reuptake inhibitor (SSRI) downregulates 5-HT7 receptors^89^, already reduced in schizophrenia.

**Figure 4.**
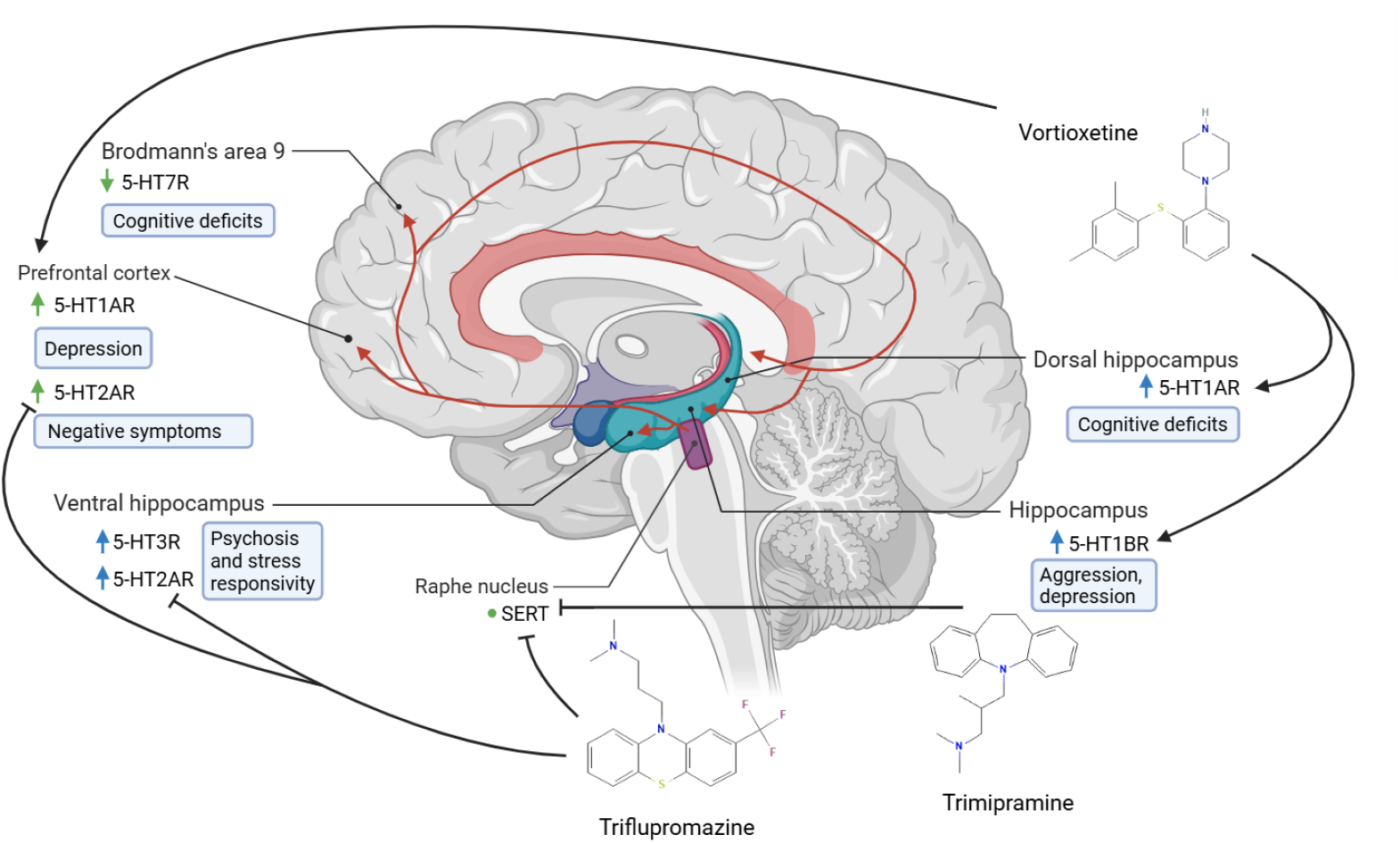
Serotonergic pathway abnormalities implicated in schizophrenia. Arrow color indicates whether a single (blue) or multiple (green) studies in the literature corroborate expression changes for the protein in schizophrenia. Black arrows signify activation, while black Ts signify inhibition. Red arrows represent serotonergic projections. Symptoms are written inside of blue rounded rectangles. 5-HT1A receptor upregulation in schizophrenia in the prefrontal cortex^95^ as well as the dorsal hippocampus is indicative of low serotonin (5-HT); in the latter region, serotonergic hypofunction is linked to cognitive deficits^96^. The 5-HT1B receptor is overexpressed within the hippocampus and its agonism has been shown to have anti-aggressive effects^97,98^. Taken together, we formulate the hypothesis that 5-HT1B overexpression is symptomatic of low 5-HT and that decreased 5-HT1B signaling causes the symptom of aggression. Prior research has shown that 5-HT1A, 5-HT1B, and 5-HT1D agonism mediates antidepressant efficacy^94^. The active conformation of the 5-HT2A receptor is upregulated in antipsychotic-free schizophrenia patients in the prefrontal cortex^99,100^. The receptor is also hyperactive in the ventral hippocampus, where it is associated with psychosis and stress responsivity, respectively. Triflupromazine, an atypical antipsychotic, antagonizes the 5-HT2A receptor and reduces negative symptoms^101^. SERT is expressed in the raphe nucleus^102,103^. Not illustrated in the figure, SERT blockers act at the axon terminals of serotonergic projections. Chronic treatment with SSRIs leads to a decrease in 5-HT2A binding potential in young subjects^104^. Increased 5-HT in the cortex as a result of SERT inhibitor administration may ameliorate 5-HT7 signaling deficits^105,106^, but further study is needed to confirm this. Overall, these findings illustrate how region-specific alterations across multiple serotonergic receptors and transporters may contribute to the cognitive, affective, and psychotic symptoms of schizophrenia.

**Vortioxetine**, a multimodal antidepressant with 5-HT1A agonism, 5-HT1B partial agonism, and 5-HT1D antagonism, improved MK-80-induced deficits in object recognition, working memory, and social behavior deficits induced by MK-801 in rats^58,59,90^. The elevated expression of 5-HT1A receptors in the prefrontal cortex and dorsal hippocampus in schizophrenia could be symptomatic of low serotonin (5-HT) in those areas (Fig 4). 5-HT1A receptor agonism counteracted phencyclidine-induced social attention deficits and has known antidepressant effects^91,92^. 5-HT1B receptor agonists had anti-aggressive effects in individuals who show moderate as well as high levels of aggression, not specific to schizophrenia^93^. A prior review suggests that activation of 5-HT1A, 5-HT1B, and 5-HT1D mediates antidepressant efficacy^94^; vortioxetine activated two of these three receptors, making it a strong candidate for treating depressive symptoms in schizophrenia.

As shown in figure 5, 5-HT1A and 5-HT7 receptors form **heterodimers**, decreasing 5-HT1A receptor-mediated activation of G(i) protein^107,108^. 5-HT1A and 5-HT2A form isoreceptor complexes, where agonism of the 5-HT2A receptor reduces affinity of a 5-HT1A agonist for its binding site^109,110^. In 5-HT transporter -/-mice, 5-HT1A receptors in the dorsal raphe nucleus show reduced binding-site density/protein levels and decreased mRNA expression, while 5-HT1B receptors in the substantia nigra show decreased binding-site density and diminished G-protein coupling^111–113^. Chronic treatment with SSRI reduces 5-HT1A-stimulated G protein activation and desensitizes terminal 5-HT1B autoreceptors^114–116^. Trimipramine and triflupromazine blocked the serotonin transporter, increasing serotonin concentration in the synaptic cleft. Effects of elevation of 5-HT include activation of the 5-HT1A receptor, 5-HT4 and 5-HT6 receptors, and 5-HT7 receptors^89,117,118^. Upon activation of the 5-HT1A receptor, inhibitory G protein beta gamma complex (Gi*βγ*) activates the phosphoinositide 3-kinase / protein kinase B (PI3K/Akt) pathway, leading to the inhibition of glycogen synthase kinase 3 (GSK3). G3K3 inhibits cyclic AMP (cAMP) response element binding protein (CREB), a transcription factor that binds to and controls the expression of brain derived neurotrophic factor (BDNF), leading to long-term neuroplastic changes and synapse regeneration^119–123^. Activation of 5-HT4 and 5-HT6 receptors stimulates the associated G*α*s subunit. This, in turn, activates the cAMP/protein kinase A (PKA) pathway, leading to the activation of extracellular signal regulated kinases 1 and 2 (ERK1/2). ERK1/2 then activates CREB, promoting the expression of (BDNF)^120,124–126^. Both activation and blockade of the 5-HT7 receptor have been shown to have beneficial effects^127^. Activation of 5-HT7, coupled to G*α*s, also leads to downstream BDNF activation^120,126^. However, activation of the 5-HT7R/MMP-9 signaling pathway has also been shown to induce depressive-like behavior, while pharmacological *blockade* of 5-HT7R produces antidepressant effects^128^. Vortioxetine activates 5-HT1AR as well as 5-HT1BR, both having been shown to mediate antidepressant efficacy^58,59,94^. These findings highlight CANDO’s utility in identifying key targets and mechanistic pathways that align with current models of schizophrenia neurobiology. The predicted drugs have the potential to address not only positive symptoms but also the negative and cognitive domains. The context-dependent therapeutic effects of both 5-HT7R activation and blockade underscore the complex pharmacodynamics of schizophrenia and reflect CANDO’s strength in capturing such multifaceted interactions. Moreover, the implicated targets suggest opportunities for modulating neural circuits disrupted in schizophrenia, offering a systems-level approach to treatment.

**Figure 5.**
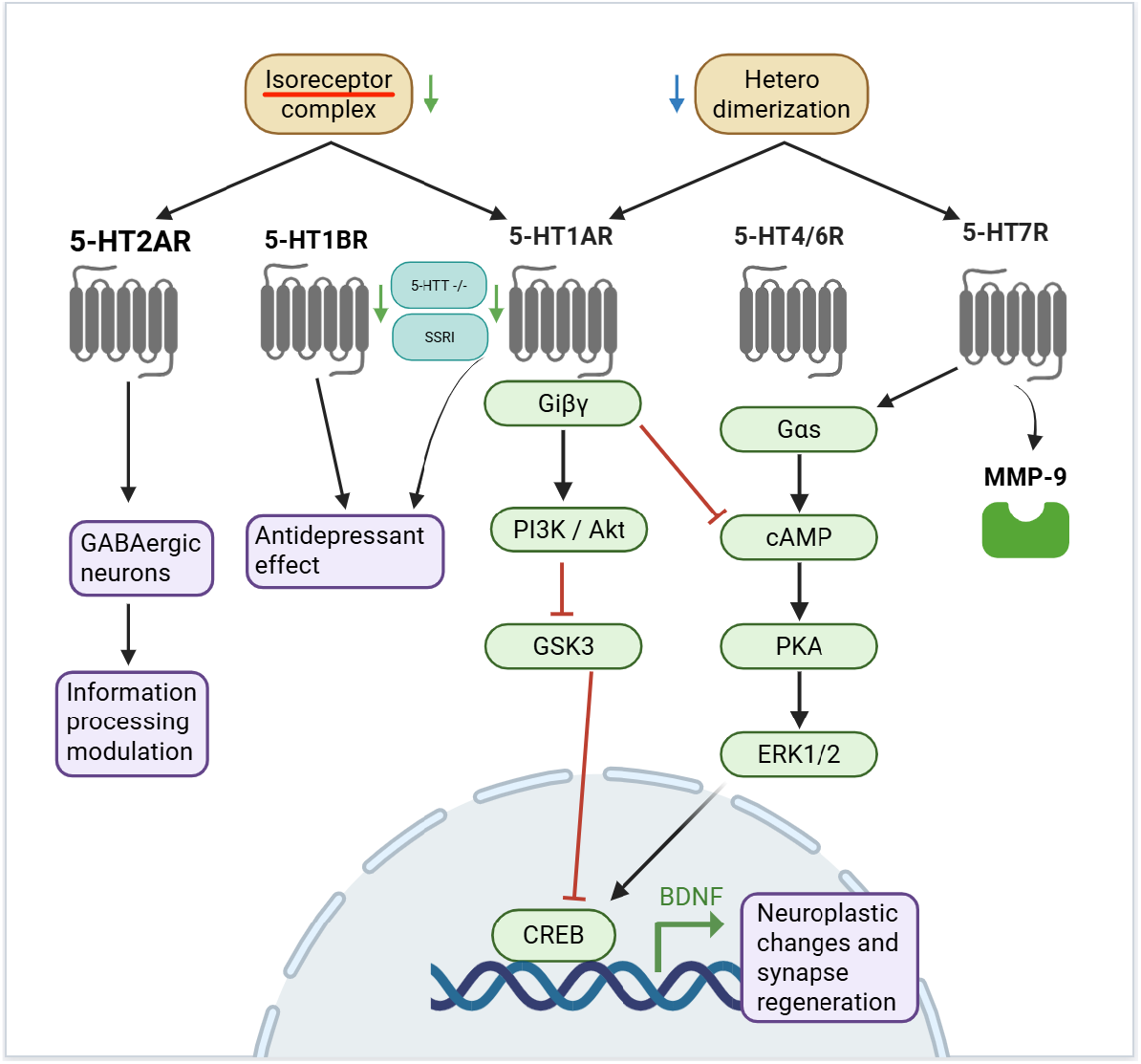
Key protein-protein interactions and downstream pathways of 5-HT receptors involved in schizophrenia pathology and treatment. Green arrows indicate a protein expression level change in schizophrenia corroborated by 2+ pieces of evidence. Blue arrows represent changes supported by a single evidence. Gold ovals refer to protein-protein interactions. Dark gray transmembrane proteins are 5-HT receptors. Green ovals are downstream proteins or subunits. Purple ovals represent physiological effects. Black arrows represent activation, while red T-shapes represent inhibition. CANDO-predicted targets map onto serotonergic mechanisms that regulate neuroplasticity and mood. Heterodimerization of 5-HT1A and 5-HT7 receptors attenuates Gi-mediated signaling, while 5-HT1A/5-HT2A isoreceptor complexes modulate receptor affinity. Agonism of 5-HT1A activates the PI3K/Akt–GSK3–CREB–BDNF pathway, promoting neuroplasticity and synapse regeneration. In contrast, 5-HT4/6 and 5-HT7 receptors signal through G*α*s–cAMP–PKA–ERK1/2–CREB cascades. Activation of the 5-HT7R/MMP-9 signaling pathway has also been shown to induce depressive like behavior, while pharmacological blockade of 5-HT7R produces antidepressant effects^128^. These pathways integrate molecular effects of predicted drugs with current neurobiological models of schizophrenia.

**Figure 6.**
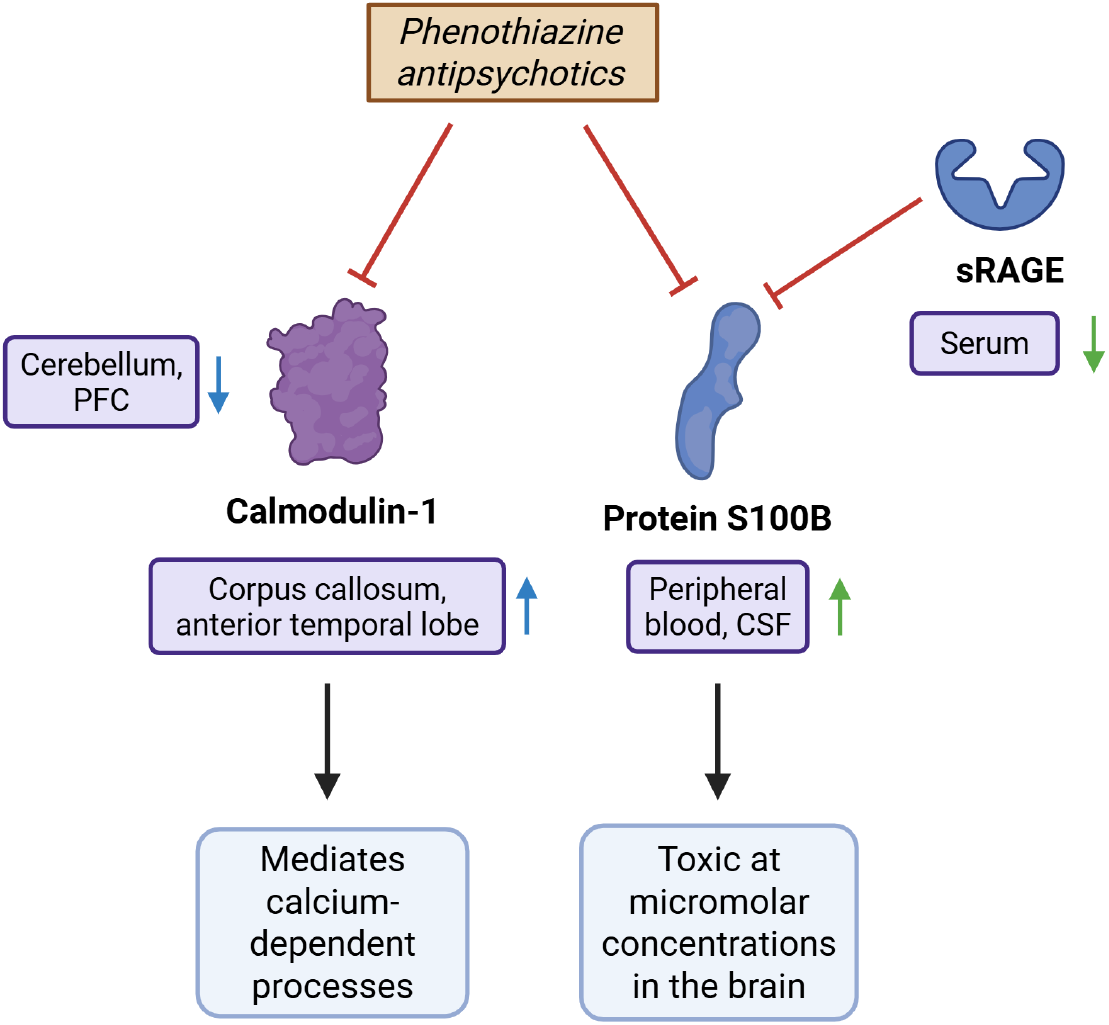
Corroborated phenothiazine antipsychotic interactions with potential therapeutic targets. Green arrows indicate a protein expression level change in schizophrenia corroborated by 2+ pieces of evidence. Blue arrows represent changes supported by a single evidence. Black arrows represent activation, while red T-shapes represent inhibition. Purple ovals indicate the locations of the expression change. Blue ovals represent physiological effects. We investigated the effect pathways of two key interactions involving calmodulin-1, the fourth-ranked consensus target prediction in our phenothiazine set, and protein S100B, the twelfth-ranked target. See Supplementary Table 3. Dysregulation of calmodulin disrupts signaling cascades and contributes to connectivity deficits in schizophrenia^73^. Calmodulin is decreased in the cerebellum, and a loss of its active form is found in the prefrontal cortex^73,135^. It is also upregulated in the corpus callosum and anterior temporal lobe in schizophrenia, but it is unknown whether this is a cause or consequence of the disorder^73^. S100B had trophic effects at nanomolar concentrations and low cellular expression of RAGE but may become toxic under conditions of micromolar concentrations, high RAGE expression, or a combination of both^74^. Increased levels of it were found in the cerebrospinal fluid and serum in schizophrenia patients^136^, while sRAGE, which sequesters and neutralizes S100B and reduces RAGE signaling, is found in reduced levels in serum^137,138^. Corroboration of the CANDO predicted protein targets with literature evidence supports the accuracy and validity of predictions. Overall, these corroborated interactions suggest that phenothiazine antipsychotics may influence calcium-dependent signaling and RAGE–related molecular pathways through modulation of calmodulin and S100B. Created in BioRender. Hu, Y. (2025) https://BioRender.com/4eued7m

#### Phenothiazine antipsychotics and calcium-binding proteins

Thiethylperazine, our fourth-ranked drug prediction with a consensus score of seven and an average rank of approximately five, is a phenothiazine atypical antipsychotic with a known interaction with the dopamine D2 receptor^129^. However, our platform ranks this interaction relatively low, 1,609 out of 20,295 proteins, with an drug-target interaction likelihood score of 0.262, substantially lower than expected for a canonical target. We observed this trend across other antipsychotics in our high-corroboration drug predictions, as only one, amisulpride, has a top corroborated interaction with dopamine receptors in Table 2. Similarly, the predicted interaction of thiethylperazine with the serotonin 5-HT2A receptor, another key target of atypical antipsychotics, is ranked 3,532 with a score of 0.193. A potential explanation for the broad side effect profiles of antipsychotics may be the ranking of high likelihood drug-protein interactions relative to known associations. We investigated these unexpected rankings by generating a phenothiazine antipsychotic consensus ranking and found that 44 of the top 50 proteins were calcium binding proteins. For comparison, calcium binding proteins comprise 39 of the consensus top 50 targets for non-phenothiazine antipsychotics, 11 for non-antipsychotic schizophrenia predictions, and none in a random set, suggesting a notable enrichment of calcium binding protein interactions among antipsychotic drugs.

#### Trifluoperazine interaction with calmodulin

The preceding observation was corroborated by existing literature reporting that neuroleptic drugs bound calmodulin, as do tricyclic antidepressants (TCAs), *in vitro*^78,79^. TCAs are present in our set of non-antipsychotic schizophrenia predictions. Recent evidence suggested that trifluoperazine binds to calmodulin-2 *in vivo* in the brain, but further studies are needed to substantiate this interaction^130^. The top ligand driving high-scoring (0.65–0.71) predicted interactions in our phenothiazines set is a calmodulin–trifluoperazine complex^131,132^. This complex has been experimentally shown to be inhibitory: trifluoperazine competes with target peptides for calmodulin binding and induces conformational changes that block calmodulin’s interaction with those targets^76,77^. Importantly, calmodulin activates Ca^2+^/calmodulin-dependent protein kinase kinase 2 (CaMKK2), and reduced CaMKK2 activity has been associated with schizophrenia in human studies^133^. As a result, inhibition of calmodulin may have a negative effect on disease prognosis. Calcium homeostasis is tightly regulated in the brain and plays a critical role in intracellular signaling cascades, including those downstream of 5-HT2A receptor activation that converge on intracellular calcium release. Calmodulin acts as the primary calcium sensor of the brain and mediates calcium-dependent processes; dysregulation may disrupt protein interactions and signaling cascades essential for cerebellar circuit formation and function. Such disruptions could contribute to the connectivity deficits observed in schizophrenia, supporting calmodulin’s potential as a therapeutic target^73^. The calmodulin-trifluoperazine complex also drove our predictions of phenothiazine interactions with several calmodulin-like proteins and myosin light chains and polypeptides. The relevance of these targets to schizophrenia remains to be elucidated, and future research may determine whether their inhibition contributes to therapeutic efficacy or to adverse effects.

#### Interaction between phenothiazines and S100B

Several S100 family proteins also appear among the top predicted targets for phenothiazines. These predictions are driven by the structural complex of S100A4 bound to a phenothiazine, and prior work has shown that chlorpromazine inhibits S100 proteins *in vitro*^80,81,134^. Previous studies indicate that S100B is a potential therapeutic target for various neural disorders, including schizophrenia, as its overexpression or administration induces disease worsening, while its deletion or inhibition produces beneficial effects^75^. Similarly, Sorci et al. report that the effects of S100B are context dependent. At nanomolar concentrations and low cellular expression of the receptor for advanced glycation end products (RAGE), S100B may have trophic effects, while at micromolar concentrations and / or in the presence of high RAGE density, it may become toxic, contributing to inflammation and cell death^74^. The protein has been shown to be upregulated in the frontal cortex and downregulated in the deep white matter of the anterior cingulate gyrus and the corpus callosum of schizophrenia patients^75^. Future research should evaluate whether inhibition of S100B can produce therapeutic benefits in schizophrenia, particularly in patient subgroups characterized by elevated S100B levels and / or high RAGE expression, and experimentally verify whether phenothiazines bind S100B *in vivo*.

#### MAO-B

High platelet levels of monoamine oxidase B (MAO-B) activity have been found to be associated with severe agitation in subjects with schizophrenia^139^. Pargyline, a MAO-B inhibitor, warrants investigation as a treatment for agitation. As an adjunctive treatment, pargyline is effective in alleviating depression, anergy, aberrant affect, mood, and self-confidence^140,141^.

#### Drugs alleviating secondary symptoms

Certain drugs in our prediction set act on pathways beyond those underlying mechanisms of delusions, hallucinations, and disorganized speech, the three positive symptoms required for a diagnosis of schizophrenia, according to the Diagnostic and Statistical Manual of Mental Disorders, Fifth Edition^142^. In most patients, these symptoms are attributed to hyperactive dopaminergic signaling in the associative striatum. Antipsychotics mitigate this dysfunction by antagonizing downstream D2 receptors, thereby reducing the severity of positive symptoms^143^. However, a subset of our predicted drugs target negative or cognitive symptoms, and these may not be part of the primary diagnostic criteria. This pattern reflects the diversity of drugs associated with schizophrenia in our library, the off-target application of CANDO, and its accuracy in predicting drugs and proteins in the process of discovering novel therapeutics. These drugs may possess polypharmacological profiles affecting multiple neurotransmitter systems. For example, the drug pizotifen, an alleviator of depressive symptoms in patients with schizophrenia, appeared across the similarity lists of several associated antipsychotic drugs that were not treatments for secondary symptoms themselves. These antipsychotics shared broad drug-proteome interaction signature overlaps with pizotifen, particularly involving the 5-HT receptor family, adrenergic receptors, and muscarinic acetylcholine receptors^144–147^. Halazepam, a drug that has been reported to reduce anxiety and tension in schizophrenia patients, was included in the similarity lists of associated drugs also used to treat anxiety^148,149^, rather than psychosis. These drugs exhibited interaction similarities with halazepam at GABA(A) receptors, the GABA(A) benzodiazepine binding site, and the translocator protein^150–154^. This limitation may, in another view, represent an advantage of the study, potentially facilitating the serendipitous discovery of therapeutics that address unmet needs or reveal novel pathways to treating certain symptoms.

#### Exclusion of investigational compounds

We used the library of approved drugs to prioritize drugs with established safety profiles for drug prediction. However, this choice excluded investigational compounds with potentially high therapeutic value.

#### Approved abroad and discontinued drugs

Our drug-indication mapping excludes drugs not approved for schizophrenia in the U.S. but are associated with the indication in other regions such as Canada and Europe. Notable among these is our best-ranking prediction, zuclopenthixol^37^. Additionally, drugs that were once approved for schizophrenia but later discontinued for reasons unrelated to efficacy or side effects, including butaperazine and perazine^39,155,156^ were also excluded. Although these drugs could have been classified as schizophrenia associated in our drug-indication mapping, it is validating that CANDO recovered them in strong positions within our predictions.

#### Additional limitations

In generating the protein target prediction lists for each drug, we used the top 100 highest-ranking proteins without applying a score cutoff, potentially including weak or excluding strong interactions. Our interaction scoring protocol considers only protein structure and not the physiological or biochemical context where a protein operates *in vivo*; as a result, actual relative binding affinities may differ from our predicted rankings. Future research will incorporate additional data that reflects the *in vivo* state of the protein, such as tissue-specific expression levels or protein conformational dynamics under physiological conditions.

#### Future directions

Future studies could expand the search space by incorporating both approved drugs and investigational compounds, as well as virtual libraries of computationally designed molecules. These compounds can be computationally screened, with candidates demonstrating the most favorable interaction profiles selected for synthesis and experimental validation^13^. Additionally, future research could refine the schizophrenia indication in CANDO by subdividing it into symptom-or domain-specific subindications and clustering drugs accordingly. These clusters could be annotated using the NIMH Research Domain Criteria (RDoC) framework, as it categorizes symptoms along dimensions such as cognition, social processes, and arousal. This stratified approach would allow CANDO to generate more precise and clinically relevant drug predictions by capturing the heterogeneity of schizophrenia. It may also enhance benchmarking performance and increase the interpretability of overlap analyses by aligning predicted targets with distinct neurobiological domains.

In summary, this study applied the multitarget CANDO platform to identify and mechanistically contextualize repurposable therapeutics for schizophrenia. Benchmarking demonstrated that CANDO reliably recaptured known drug–indication relationships and prioritized clinically relevant compounds, while literature review corroborated 25 high-confidence candidates spanning phenothiazines, benzodiazepines, TCAs, and MAO inhibitors. Overlap analyses revealed that predicted protein targets align with established schizophrenia biology, with targets of our top-ranked drug predictions enriched for canonical neurotransmitter receptors, monoamine transporters, and calcium-binding proteins. These findings support the utility of a proteome-wide approach for uncovering both known and novel therapeutic mechanisms, and highlight calmodulin- and S100-related signaling as potential avenues for future experimental validation. Together, this work demonstrates the utility of CANDO for prioritizing repurposable drug candidates for schizophrenia and identifying the protein targets and pathways most likely to mediate their effects. These candidates and associated target pathways provide a concrete starting point for experimental testing and for advancing polypharmacological approaches to schizophrenia therapeutics.

## Author contributions statement

Y.H. performed the experiments and carried out the data analysis. Y.H., Z.F., and R.S. jointly contributed to the experimental design and interpretation. Y.H. drafted the initial manuscript, and all authors reviewed, edited, and approved the final version. R.S. supervised the project.”

## Funding statement

This study was funded through a National Institutes of Health (NIH) Director’s Pioneer Award (DP1OD006779), a NIH Clinical and Translational Sciences (NCATS) Award (UL1TR001412), a NIH National Library of Medicine (NLM) T15 Award (T15LM012495), a NIH NLM R25 Award (R25LM014213), a NIH NCATS ASPIRE Design Challenge Award, a NIH NCATS ASPIRE Reduction-to-Practice Award, a National Institute of Standards of Technology (NIST) Award (60NANB22D168), a NIDA Mentored Research Scientist Development Award (K01DA056690), and startup funds from the Department of Biomedical Informatics at the University at Buffalo. Disclaimer: The content is solely the responsibility of the authors and does not necessarily represent the official views of the National Institutes of Health.

## Additional information

The authors disclose that they formed multiple startup ventures engaged in the commercialization of outputs derived from the CANDO platform. The funders had no involvement in the study’s conception or design, the acquisition, analysis, or interpretation of data, the preparation of the manuscript, or the decision to submit it for publication.

## Notes

### Summary of Updates

The manuscript has undergone several rounds of internal revision that have substantially improved it. We have also added a funding statement to the PDF.

## References

1. Kahn, R. S. et al. Schizophrenia. Nat. Rev. Dis. Primers 1, 15067, DOI: 10.1038/nrdp.2015.67 (2015).

2. National Institute of Mental Health. Schizophrenia (2022).

3. Patel, K. R., Cherian, J., Gohil, K. & Atkinson, D. Schizophrenia: Overview and treatment options. Pharm. Ther. 39, 638–645 (2014).

4. Zeng, G. et al. Factors affecting negative symptoms in schizophrenia and their relationship with anxiety and depression. Neuropsychiatr. Dis. Treat. 21, 229–240, DOI: 10.2147/NDT.S492849 (2025).

5. Mizuno, Y., McCutcheon, R. A., Brugger, S. P. & Howes, O. D. Heterogeneity and efficacy of antipsychotic treatment for schizophrenia with or without treatment resistance: a meta-analysis. Neuropsychopharmacology 45, 622–631, DOI: 10.1038/s41386-019-0577-3 (2020).

6. World Health Organization. Schizophrenia fact sheet (2022).

7. Lee, R. et al. Predicting treatment resistance in positive and negative symptom domains from first episode psychosis: Development of a clinical prediction model. Schizophr. Res. 274, 66–77, DOI: 10.1016/j.schres.2024.09.010 (2024).

8. Meltzer, H. Y. & Gadaleta, E. Contrasting typical and atypical antipsychotic drugs. Focus 19, 3–13, DOI: 10.1176/appi.focus.20200051 (2021).

9. Brisch, R. et al. The role of dopamine in schizophrenia from a neurobiological and evolutionary perspective: old fashioned, but still in vogue. Front. Psychiatry 5, 47, DOI: 10.3389/fpsyt.2014.00047 (2014). Erratum in: Front Psychiatry. 2014;5:110. Author corrections: Braun, Katharina; Kumaratilake, Jaliya.

10. Reimers, A., Odin, P. & Ljung, H. Drug-induced cognitive impairment. Drug Saf. 48, 339–361, DOI: 10.1007/s40264-024-01506-5 (2025).

11. Lin, X., Li, X. & Lin, X. A review on applications of computational methods in drug screening and design. Mol. (Basel, Switzerland) 25, 1375, DOI: 10.3390/molecules25061375 (2020).

12. Sadybekov, A. V. & Katritch, V. Computational approaches streamlining drug discovery. Nature 616, 673–685, DOI: 10.1038/s41586-023-05905-z (2023).

13. Oliveira, T. A. d., Silva, M. P. d., Maia, E. H. B., Silva, A. M. d. & Taranto, A. G. Virtual screening algorithms in drug discovery: A review focused on machine and deep learning methods. Drugs Drug Candidates 2, 311–334, DOI: 10.3390/ddc2020017 (2023).

14. Vázquez, J., López, M., Gibert, E., Herrero, E. & Luque, F. J. Merging ligand-based and structure-based methods in drug discovery: An overview of combined virtual screening approaches. Mol. (Basel, Switzerland) 25, 4723, DOI: 10.3390/molecules25204723 (2020).

15. Abramson, J., Adler, J., Dunger, J. et al. Accurate structure prediction of biomolecular interactions with alphafold 3. Nature 630, 493–500, DOI: 10.1038/s41586-024-07487-w (2024).

16. Fine, J., Lackner, R., Samudrala, R. et al. Computational chemoproteomics to understand the role of selected psychoactives in treating mental health indications. Sci. Reports 9, 13155, DOI: 10.1038/s41598-019-49515-0 (2019).

17. Minie, M. et al. Cando and the infinite drug discovery frontier. Drug Discov. Today 19, 1353–1363, DOI: 10.1016/j.drudis.2014.06.018 (2014).

18. Mangione, W., Falls, Z., Melendy, T., Chopra, G. & Samudrala, R. Shotgun drug repurposing biotechnology to tackle epidemics and pandemics. Drug Discov. Today 25, 1126–1128, DOI: 10.1016/j.drudis.2020.05.002 (2020).

19. Falls, Z., Mangione, W., Schuler, J. & Samudrala, R. Exploration of interaction scoring criteria in the cando platform. BMC Res. Notes 12, 318, DOI: 10.1186/s13104-019-4356-3 (2019).

20. Mangione, W., Falls, Z. & Samudrala, R. Optimal covid-19 therapeutic candidate discovery using the cando platform. Front. Pharmacol. 13, 1031689, DOI: 10.3389/fphar.2022.970494 (2022).

21. Jenwitheesuk, E., Horst, J. A., Rivas, K. L., Van Voorhis, W. C. & Samudrala, R. Novel paradigms for drug discovery: computational multitarget screening. Trends Pharmacol. Sci. 29, 62–71, DOI: 10.1016/j.tips.2007.11.007 (2008).

22. Chopra, G., Kaushik, S., Elkin, P. L. & Samudrala, R. Combating ebola with repurposed therapeutics using the cando platform. Mol. (Basel, Switzerland) 21, 1537, DOI: 10.3390/molecules21121537 (2016).

23. Bruggemann, L. et al. Multiscale analysis and validation of effective drug combinations targeting driver kras mutations in non-small cell lung cancer. Int. J. Mol. Sci. 24, 997, DOI: 10.3390/ijms24020997 (2023).

24. Palanikumar, L. et al. Protein mimetic amyloid inhibitor potently abrogates cancer-associated mutant p53 aggregation and restores tumor suppressor function. Nat. Commun. 12, 3962, DOI: 10.1038/s41467-021-23985-1 (2021).

25. Xu, S. et al. Multiscale analysis and optimal glioma therapeutic candidate discovery using the cando platform. bioRxiv 2025.05.19.654757, DOI: 10.1101/2025.05.19.654757 (2025). Preprint.

26. Davis, A. P. et al. Comparative toxicogenomics database (ctd): update 2021. Nucleic Acids Res. 49, D1138–D1143, DOI: 10.1093/nar/gkaa891 (2021).

27. Lipscomb, C. E. Medical subject headings (mesh). Bull. Med. Libr. Assoc. 88, 265–266 (2000).

28. Mammen, M. J. et al. Proteomic network analysis of bronchoalveolar lavage fluid in ex-smokers to discover implicated protein targets and novel drug treatments for chronic obstructive pulmonary disease. Pharm. (Basel, Switzerland) 15, 566, DOI: 10.3390/ph15050566 (2022).

29. Moukheiber, L. et al. Identifying protein features and pathways responsible for toxicity using machine learning and tox21: Implications for predictive toxicology. Mol. (Basel, Switzerland) 27, 3021, DOI: 10.3390/molecules27093021 (2022).

30. Schuler, J. et al. Evaluating the performance of drug-repurposing technologies. Drug Discov. Today 27, 49–64, DOI: 10.1016/j.drudis.2021.08.002 (2022).

31. Van Norden, M., Mangione, W., Falls, Z. & Samudrala, R. Strategies for robust, accurate, and generalizable benchmarking of drug discovery platforms. bioRxiv DOI: 10.1101/2024.12.10.627863 (2024). Preprint.

32. Järvelin, K. & Kekäläinen, J. Cumulated gain-based evaluation of ir techniques. ACM Transactions on Inf. Syst. 20, 422–446, DOI: 10.1145/582415.582418 (2002).

33. Gao, J. Identification of Putative Drug Candidates for Type 2 Diabetes Using the CANDO Platform. Master’s thesis, University at Buffalo, The State University of New York, Buffalo, NY (2025). A thesis submitted to the Faculty of the Graduate School.

34. Jaccard, P. The distribution of the flora in the alpine zone. 1. New Phytol. 11, 37–50 (1912).

35. Chung, N. C., Miasojedow, B., Startek, M. & Gambin, A. Jaccard/tanimoto similarity test and estimation methods for biological presence-absence data. BMC Bioinforma. 20, 644, DOI: 10.1186/s12859-019-3118-5 (2019).

36. U.S. Food and Drug Administration, Center for Drug Evaluation and Research. Orange book preface to the 45th edition — approved drug products with therapeutic equivalence evaluations. https://www.fda.gov/drugs/development-approval-process-drugs/orange-book-preface (2024). Accessed: 2025-11-01.

37. DrugBank. Zuclopenthixol. https://go.drugbank.com/drugs/DB01624 (Accessed 2025). DrugBank ID: DB01624.

38. DrugBank. Butaperazine. https://go.drugbank.com/drugs/DB13213 (Accessed 2025). DrugBank ID: DB13213.

39. DrugBank. Perazine. https://go.drugbank.com/drugs/DB12710 (Accessed 2025). DrugBank ID: DB12710.

40. DrugBank. Lumateperone. https://go.drugbank.com/drugs/DB06077 (Accessed 2025). DrugBank ID: DB06077.

41. DrugBank. Prochlorperazine. https://go.drugbank.com/drugs/DB00433 (Accessed 2025). DrugBank ID: DB00433.

42. Haddad, P., Taylor, M., Patel, M. X. & Taylor, D. Guidance on switching away from piportil depot® (pipotiazine palmitate) injection. Br. J. Psychiatry 206, 521–521, DOI: 10.1192/bjp.206.6.521 (2015).

43. DrugBank. Promazine. https://go.drugbank.com/drugs/DB00420 (Accessed 2025). DrugBank ID: DB00420.

44. DrugBank. Clomipramine. https://go.drugbank.com/drugs/DB01242 (Accessed 2025). DrugBank ID: DB01242.

45. DrugBank. Sultopride. https://go.drugbank.com/drugs/DB13273 (Accessed 2025). DrugBank ID: DB13273.

46. U.S. Food and Drug Administration. Approved drug products with therapeutic equivalence evaluations (orange book). https://www.accessdata.fda.gov/scripts/cder/ob/ (2025). Accessed: 2025-12-10.

47. DrugBank. Amisulpride. https://go.drugbank.com/drugs/DB06288 (Accessed 2025). DrugBank ID: DB06288.

48. U.S. Food and Drug Administration. Label for nda 209510 — prescribing information. https://www.accessdata.fda.gov/drugsatfda_docs/label/2020/209510s000lbl.pdf (2020). Accessed: 2025-12-10.

49. U.S. Food and Drug Administration. Vesprin (triflupromazine hydrochloride). FDA Orange Book: Approved Drug Products with Therapeutic Equivalence Evaluations (1982). NDA 011123; Proprietary name: VESPRIN; Active ingredient: Triflupromazine Hydrochloride; Applicant holder: Bristol Myers Squibb Co.; Approval date: Prior to Jan 1, 1982.

50. DrugBank. Triflupromazine. https://go.drugbank.com/drugs/DB00508 (Accessed 2025). DrugBank ID: DB00508.

51. DrugBank. lurasidone. https://go.drugbank.com/drugs/DB08815 (Accessed 2025). DrugBank ID: DB08815.

52. Ouyang, X. et al. Rapid screening of acetylcholinesterase inhibitors by effect-directed analysis using LC × LC fractionation, a high throughput in vitro assay, and parallel identification by time of flight mass spectrometry. Anal. Chem. 88, 2353–2360, DOI: 10.1021/acs.analchem.5b04311 (2016).

53. DrugBank. Phenelzine. https://go.drugbank.com/drugs/DB00780 (Accessed 2025). DrugBank ID: DB00780.

54. Malizia, A. L. et al. Demonstration of clomipramine and venlafaxine occupation at serotonin reuptake sites in man in vivo. J. Psychopharmacol. 11, 279–281, DOI: 10.1177/026988119701100312 (1997).

55. Gerile, S. C. et al. Inhibitory action of antidepressants on mouse betaine/gaba transporter (bgt1) heterologously expressed in cell cultures. Int. J. Mol. Sci. 13, 2578–2589, DOI: 10.3390/ijms13032578 (2012). Epub 2012 Feb 24.

56. Tatsumi, M., Jansen, K., Blakely, R. D. & Richelson, E. Pharmacological profile of neuroleptics at human monoamine transporters. Eur. J. Pharmacol. 368, 277–283, DOI: 10.1016/s0014-2999(99)00005-9 (1999).

57. DrugBank. Trimipramine. https://go.drugbank.com/drugs/DB00726 (Accessed 2025). DrugBank ID: DB00726.

58. DrugBank. Vortioxetine. https://go.drugbank.com/drugs/DB09068 (Accessed 2025). DrugBank ID: DB09068.

59. Sanchez, C., Asin, K. E. & Artigas, F. Vortioxetine, a novel antidepressant with multimodal activity: Review of preclinical and clinical data. Pharmacol. & Ther. 145, 43–57, DOI: 10.1016/j.pharmthera.2014.07.001 (2015).

60. Chakroborty, S. et al. Impact of vortioxetine on synaptic integration in prefrontal-subcortical circuits: Comparisons with escitalopram. Front. Pharmacol. 8, 764, DOI: 10.3389/fphar.2017.00764 (2017).

61. Tollens, F. et al. The affinity of antipsychotic drugs to dopamine and serotonin 5-ht2 receptors determines their effects on prefrontal-striatal functional connectivity. Eur. Neuropsychopharmacol. 28, 1035–1046, DOI: 10.1016/j.euroneuro.2018.05.016 (2018).

62. Donahue, T. J. et al. Examination of the mechanisms underlying the discriminative stimulus properties of the atypical antipsychotic amisulpride. Behav. Pharmacol. 35, 47–54, DOI: 10.1097/FBP.0000000000000760 (2024). Epub 2023 Nov 15.

63. Meltzer, H. Y. Serotonergic mechanisms as targets for existing and novel antipsychotics. In Handbook of Experimental Pharmacology, 212, 87–124, DOI: 10.1007/978-3-642-25761-2_4 (Springer Science and Business Media Deutschland GmbH, 2012).

64. Meltzer, H. Y. & Massey, B. W. The role of serotonin receptors in the action of atypical antipsychotic drugs. Curr. Opin. Pharmacol. 11, 59–67, DOI: 10.1016/j.coph.2011.02.007 (2011). Epub 2011 Mar 21.

65. Castelli, M. P., Mocci, I., Sanna, A. M., Gessa, G. L. & Pani, L. (-)s amisulpride binds with high affinity to cloned dopamine d3 and d2 receptors. Eur. J. Pharmacol. 432, 143–147, DOI: 10.1016/s0014-2999(01)01484-4 (2001).

66. Vasse, M. & Protais, P. Increased grooming behaviour is induced by apomorphine in mice treated with discriminant benzamide derivatives. Eur. J. Pharmacol. 156, 1–11, DOI: 10.1016/0014-2999(88)90141-0 (1988).

67. Chivers, J. K., Gommeren, W., Leysen, J. E., Jenner, P. & Marsden, C. D. Comparison of the in-vitro receptor selectivity of substituted benzamide drugs for brain neurotransmitter receptors. J. Pharm. Pharmacol. 40, 415–421, DOI: 10.1111/j.2042-7158.1988.tb06306.x (1988).

68. Sahyoun, H. A., Costall, B. & Naylor, R. J. Benzamide action at α_2_-adrenoceptors modifies catecholamine-induced contraction and relaxation of circular smooth muscle from guinea-pig stomach. Naunyn-Schmiedeberg’s Arch. Pharmacol. 319, 8–11, DOI: 10.1007/BF00491470 (1982).

69. DrugBank. Flunarizine. https://go.drugbank.com/drugs/DB04841 (Accessed 2025). DrugBank ID: DB04841.

70. Takao, K. et al. 2-styrylchromone derivatives as potent and selective monoamine oxidase b inhibitors. Bioorganic Chem. 92, 103285, DOI: 10.1016/j.bioorg.2019.103285 (2019). Epub 2019 Sep 18.

71. Dyck, L. E. & Dewar, K. M. Inhibition of aromatic l-amino acid decarboxylase and tyrosine aminotransferase by the monoamine oxidase inhibitor phenelzine. J. Neurochem. 46, 1899–1903, DOI: 10.1111/j.1471-4159.1986.tb08511.x (1986).

72. Tan, A. H. Y. et al. Lysine-specific histone demethylase 1a regulates macrophage polarization and checkpoint molecules in the tumor microenvironment of triple-negative breast cancer. Front. Immunol. 10, 1351, DOI: 10.3389/fimmu.2019.01351 (2019).

73. Vidal-Domènech, F. et al. Calcium-binding proteins are altered in the cerebellum in schizophrenia. PLOS ONE 15, e0230400, DOI: 10.1371/journal.pone.0230400 (2020).

74. Sorci, G. et al. S100b protein, a damage-associated molecular pattern protein in the brain and heart, and beyond. Cardiovasc. Psychiatry Neurol. 2010, 656481, DOI: 10.1155/2010/656481 (2010).

75. Michetti, F. et al. The s100b story: from biomarker to active factor in neural injury. J. Neurochem. 148, 168–187, DOI: 10.1111/jnc.14574 (2018).

76. Vertessy, B. G. et al. Simultaneous binding of drugs with different chemical structures to ca2+-calmodulin: Crystallographic and spectroscopic studies. Biochemistry 37, 15300–15310, DOI: 10.1021/bi980795a (1998). PMID: 9799490.

77. Vandonselaar, M., Hickie, R. A., Quail, J. W. & Delbaere, L. T. J. Trifluoperazine-induced conformational change in ca2+-calmodulin. Nat. Struct. Biol. 1, 795–801, DOI: 10.1038/nsb1194-795 (1994).

78. Lejoyeux, M. & Maziere, J. C. Could the interaction of neuroleptics with calmodulin be an “explanation” of the psychotropic effects? Encephale 17, 11–15 (1991).

79. Roufogalis, B. D., Minocherhomjee, A. M. & Al-Jobore, A. Pharmacological antagonism of calmodulin. Can. J. Biochem. Cell Biol. 61, 927–933, DOI: 10.1139/o83-118 (1983).

80. Donato, R. Chlorpromazine inhibits the calcium-mediated effects of s-100 protein(s) on assembled brain microtubule proteins, but not those on microtubule protein assembly. Biochem. Biophys. Res. Commun. 122, 983–990, DOI: 10.1016/0006-291X(84)91188-4 (1984).

81. Malashkevich, V. N. et al. Phenothiazines inhibit s100a4 function by inducing protein oligomerization. Proc. Natl. Acad. Sci. United States Am. 107, 8605–8610, DOI: 10.1073/pnas.0913660107 (2010).

82. Gao, M. et al. Association between efhd2 gene polymorphisms and schizophrenia among the han population in northern china. J. Int. Med. Res. 48, 300060520932801, DOI: 10.1177/0300060520932801 (2020).

83. Murakami, M., Nagahama, M., Abe, Y. & Niikura, T. Humanin affects object recognition and gliosis in short-term cuprizone-treated mice. Neuropeptides 66, 90–96, DOI: 10.1016/j.npep.2017.10.002 (2017).

84. da Silva, F. E. R. et al. Sex and the estrous-cycle phase influence the expression of g protein-coupled estrogen receptor 1 (gper) in schizophrenia: Translational evidence for a new target. Mol. Neurobiol. 60, 3650–3663, DOI: 10.1007/s12035-023-03295-x (2023).

85. Barrera-Conde, M. et al. Role of cyclin-dependent kinase 5 in psychosis and the modulatory effects of cannabinoids. Neurobiol. Dis. 176, 105942, DOI: 10.1016/j.nbd.2022.105942 (2023).

86. Tatsumi, M., Jansen, K., Blakely, R. D. & Richelson, E. Pharmacological profile of neuroleptics at human monoamine transporters. Eur. J. Pharmacol. 368, 277–283, DOI: 10.1016/S0014-2999(99)00005-9 (1999).

87. Vestre, N. D., Hall, W. B. & Schiele, B. C. A comparison of fluphenazine, triflupromazine, and phenobarbital in the treatment of chronic schizophrenic patients: a double-blind controlled study. J. Clin. Exp. Psychopathol. & Q. Rev. Psychiatry Neurol. 23, 149–159 (1962).

88. Eikmeier, G. et al. Trimipramine–an atypical neuroleptic? Int. Clin. Psychopharmacol. 6, 147–153, DOI: 10.1097/00004850-199100630-00003 (1991).

89. Mullins, U., Gianutsos, G. & Eison, A. Effects of antidepressants on 5-ht7 receptor regulation in the rat hypothalamus. Neuropsychopharmacology 21, 352–367, DOI: 10.1016/S0893-133X(99)00041-X (1999).

90. Bozkurt, N. M. & Unal, G. Vortioxetine improved negative and cognitive symptoms of schizophrenia in subchronic mk-801 model in rats. Behav. Brain Res. 444, 114365, DOI: 10.1016/j.bbr.2023.114365 (2023).

91. Boulay, D. et al. Ssr181507, a putative atypical antipsychotic with dopamine d2 antagonist and 5-ht1a agonist activities: improvement of social interaction deficits induced by phencyclidine in rats. Neuropharmacology 46, 1121–1129, DOI: 10.1016/j.neuropharm.2004.02.008 (2004).

92. Kennett, G. A., Dourish, C. T. & Curzon, G. Antidepressant-like action of 5-ht1a agonists and conventional antidepressants in an animal model of depression. Eur. J. Pharmacol. 134, 265–274, DOI: 10.1016/0014-2999(87)90357-8 (1987).

93. de Almeida, R. M. M. & Miczek, K. A. Aggression escalated by social instigation or by discontinuation of reinforcement (“frustration”) in mice: Inhibition by anpirtoline: A 5-ht1b receptor agonist. Neuropsychopharmacology 27, 171–181, DOI: 10.1016/S0893-133X(02)00291-9 (2002).

94. David, D. J. & Gardier, A. M. Les bases de pharmacologie fondamentale du système sérotoninergique : application à la réponse antidépressive [the pharmacological basis of the serotonin system: Application to antidepressant response]. L’Encephale 42, 255–263, DOI: 10.1016/j.encep.2016.03.012 (2016).

95. Banerjee, P., Mehta, M. & Kanjilal, B. The 5-ht1a receptor: A signaling hub linked to emotional balance. In Chattopadhyay, A. (ed.) Serotonin Receptors in Neurobiology, chap. 7 (CRC Press/Taylor & Francis, Boca Raton (FL), 2007).

96. Kandilakis, C. L. & Papatheodoropoulos, C. Serotonin modulation of dorsoventral hippocampus in physiology and schizophrenia. Int. J. Mol. Sci. 26, 7253, DOI: 10.3390/ijms26157253 (2025).

97. de Almeida, R. M. M. & Miczek, K. A. Aggression escalated by social instigation or by discontinuation of reinforcement (“frustration”) in mice: Inhibition by anpirtoline: A 5-ht1b receptor agonist. Neuropsychopharmacology 27, 171–181, DOI: 10.1016/S0893-133X(02)00291-9 (2002).

98. López-Figueroa, A. L. et al. Serotonin 5-ht1a, 5-ht1b, and 5-ht2a receptor mrna expression in subjects with major depression, bipolar disorder, and schizophrenia. Biol. Psychiatry 55, 225–233, DOI: 10.1016/j.biopsych.2003.09.017 (2004). Erratum in: Biological Psychiatry. 2004 Mar 15;55(6):660.

99. Muguruza, C. et al. Dysregulated 5-ht(2a) receptor binding in postmortem frontal cortex of schizophrenic subjects. Eur. Neuropsychopharmacol. 23, 852–864, DOI: 10.1016/j.euroneuro.2012.10.006 (2013). Epub 2012 Nov 21.

100. Diez-Alarcia, R. et al. Opposite alterations of 5-ht2a receptor brain density in subjects with schizophrenia: relevance of radiotracers pharmacological profile. Transl. Psychiatry 11, 302, DOI: 10.1038/s41398-021-01430-7 (2021).

101. Romeo, B., Willaime, L., Rari, E., Benyamina, A. & Martelli, C. Efficacy of 5-ht2a antagonists on negative symptoms in patients with schizophrenia: A meta-analysis. Psychiatry Res. 321, 115104, DOI: 10.1016/j.psychres.2023.115104 (2023). Epub 2023 Feb 8.

102. Chen, H. T., Clark, M. & Goldman, D. Quantitative autoradiography of 3h-paroxetine binding sites in rat brain. J. Pharmacol. Toxicol. Methods 27, 209–216, DOI: 10.1016/1056-8719(92)90043-z (1992).

103. Austin, M. C., Bradley, C. C., Mann, J. J. & Blakely, R. D. Expression of serotonin transporter messenger rna in the human brain. J. Neurochem. 62, 2362–2367, DOI: 10.1046/j.1471-4159.1994.62062362.x (1994).

104. Meyer, J. H. et al. The effect of paroxetine on 5-ht(2a) receptors in depression: an [18f]setoperone pet imaging study. Am. J. Psychiatry 158, 78–85, DOI: 10.1176/appi.ajp.158.1.78 (2001).

105. Dean, B., Pavey, G., Thomas, D. & Scarr, E. Cortical serotonin7, 1d and 1f receptors: effects of schizophrenia, suicide and antipsychotic drug treatment. Schizophr. Res. 88, 265–274, DOI: 10.1016/j.schres.2006.07.003 (2006). Epub 2006 Aug 17.

106. East, S. Z., Burnet, P. W., Kerwin, R. W. & Harrison, P. J. An rt-pcr study of 5-ht6 and 5-ht7 receptor mrnas in the hippocampal formation and prefrontal cortex in schizophrenia. Schizophr. Res. 57, 15–26, DOI: 10.1016/S0920-9964(01)00323-1 (2002).

107. Renner, U. et al. Heterodimerization of serotonin receptors 5-ht1a and 5-ht7 differentially regulates receptor signalling and trafficking. J. Cell Sci. 125, 2486–2499, DOI: 10.1242/jcs.101337 (2012). Epub 2012 Feb 22.

108. Prasad, S., Ponimaskin, E. & Zeug, A. Serotonin receptor oligomerization regulates camp-based signaling. J. Cell Sci. 132, jcs230334, DOI: 10.1242/jcs.230334 (2019).

109. Borroto-Escuela, D. O. et al. Existence of brain 5-ht1a-5-ht2a isoreceptor complexes with antagonistic allosteric receptor-receptor interactions regulating 5-ht1a receptor recognition. ACS Omega 2, 4779–4789, DOI: 10.1021/acsomega.7b00629 (2017). Epub 2017 Aug 22.

110. Szlachta, M. et al. P.1.006 - chronic administration of clozapine increases the level of serotonin 5-ht1a receptor heterodimerisation with 5-ht2a or d2 receptors in the mouse cortex. Eur. Neuropsychopharmacol. 28, S7–S8, DOI: 10.1016/j.euroneuro.2017.12.024 (2018). Abstracts of the ECNP Workshop for Junior Scientists in Europe15-18 March 2017, Nice, France.

111. Fabre, V. et al. Altered expression and functions of serotonin 5-ht1a and 5-ht1b receptors in knock-out mice lacking the 5-ht transporter. Eur. J. Neurosci. 12, 2299–2310, DOI: 10.1046/j.1460-9568.2000.00126.x (2000).

112. Li, Q., Wichems, C., Heils, A., Lesch, K. P. & Murphy, D. L. Reduction in the density and expression, but not g-protein coupling, of serotonin receptors (5-ht1a) in 5-ht transporter knock-out mice: gender and brain region differences. J. Neurosci. 20, 7888–7895, DOI: 10.1523/JNEUROSCI.20-21-07888.2000 (2000).

113. Shanahan, N. A. et al. Chronic reductions in serotonin transporter function prevent 5-ht1b-induced behavioral effects in mice. Biol. Psychiatry 65, 401–408, DOI: 10.1016/j.biopsych.2008.09.026 (2009).

114. Bahri, S., Mnie-Filali, O., Dkhissi-Benyahya, O., Abrial, E. & Haddjeri, N. Neuroadaptations of the 5-ht system induced by antidepressant treatments: Old and new strategies. Annals Adv. Depress. Anxiety 1, 100001, DOI: 10.24966/AAD-7276/100001 (2014). Published Aug 18, 2014; Received Jun 19, 2014; Accepted Aug 04, 2014.

115. Pejchal, T., Foley, M. A., Kosofsky, B. E. & Waeber, C. Chronic fluoxetine treatment selectively uncouples raphe 5-ht(1a) receptors as measured by [35s]-gtpγs autoradiography. Br. J. Pharmacol. 135, 1115–1122, DOI: 10.1038/sj.bjp.0704555 (2002).

116. Shalom, G., Gur, E., Van de Kar, L. D. & Newman, M. E. Repeated administration of the 5-ht(1b) receptor antagonist sb-224289 blocks the desensitisation of 5-ht(1b) autoreceptors induced by fluoxetine in rat frontal cortex. Naunyn-Schmiedeberg’s Arch. Pharmacol. 370, 84–90, DOI: 10.1007/s00210-004-0958-x (2004).

117. Vahid-Ansari, F., Zhang, M., Zahrai, A. & Albert, P. R. Overcoming resistance to selective serotonin reuptake inhibitors: Targeting serotonin, serotonin-1a receptors and adult neuroplasticity. Front. Neurosci. 13, DOI: 10.3389/fnins.2019.00404 (2019).

118. Svenningsson, P. et al. Involvement of striatal and extrastriatal darpp-32 in biochemical and behavioral effects of fluoxetine (prozac). Proc. Natl. Acad. Sci. United States Am. 99, 3182–3187, DOI: 10.1073/pnas.052712799 (2002).

119. Hsiung, S. C., Tamir, H., Franke, T. F. & Liu, K. P. Roles of extracellular signal-regulated kinase and akt signaling in coordinating nuclear transcription factor-kappab-dependent cell survival after serotonin 1a receptor activation. J. Neurochem. 95, 1653–1666, DOI: 10.1111/j.1471-4159.2005.03496.x (2005).

120. Wong, T.-S. et al. G protein-coupled receptors in neurodegenerative diseases and psychiatric disorders. Signal Transduct. Target. Ther. 8, 177, DOI: 10.1038/s41392-023-01427-2 (2023).

121. Polter, A. M. & Li, X. 5-ht1a receptor-regulated signal transduction pathways in brain. Cell. Signal. 22, 1406–1412, DOI: 10.1016/j.cellsig.2010.03.019 (2010).

122. Xu, T., Finkbeiner, S., Arnold, D. B., Shaywitz, A. J. & Greenberg, M. E. Ca2+ influx regulates bdnf transcription by a creb family transcription factor-dependent mechanism. Neuron 20, 709–726, DOI: 10.1016/S0896-6273(00)81010-7 (1998).

123. Björkholm, C. & Monteggia, L. M. Bdnf – a key transducer of antidepressant effects. Neuropharmacology 102, 72–79, DOI: 10.1016/j.neuropharm.2015.10.034 (2016).

124. Galligan, J. J. Colonic 5-ht4 receptors are targets for novel prokinetic drugs. Neurogastroenterol. Motil. 33, e14125, DOI: 10.1111/nmo.14125 (2021).

125. Deraredj Nadim, W. et al. Physical interaction between neurofibromin and serotonin 5-ht6 receptor promotes receptor constitutive activity. Proc. Natl. Acad. Sci. United States Am. 113, 12310–12315, DOI: 10.1073/pnas.1600914113 (2016).

126. Norum, J. H., Hart, K. & Levy, F. O. Ras-dependent erk activation by the human gs-coupled serotonin receptors 5-ht4(b) and 5-ht7(a). J. Biol. Chem. 278, 3098–3104, DOI: 10.1074/jbc.M206237200 (2003).

127. Nikiforuk, A. Targeting the serotonin 5-ht7 receptor in the search for treatments for cns disorders: Rationale and progress to date. CNS Drugs 29, 265–275, DOI: 10.1007/s40263-015-0236-0 (2015).

128. Bijata, M. et al. Activation of the 5-ht7 receptor and mmp-9 signaling module in the hippocampal ca1 region is necessary for the development of depressive-like behavior. Cell Reports 38, 110532, DOI: 10.1016/j.celrep.2022.110532 (2022).

129. DrugBank. Thiethylperazine. https://go.drugbank.com/drugs/DB00372 (Accessed 2025). DrugBank ID: DB00372.

130. Kang, S. et al. Trifluoperazine, a well-known antipsychotic, inhibits glioblastoma invasion by binding to calmodulin and disinhibiting calcium release channel ip3r. Mol. Cancer Ther. 16, 217–227, DOI: 10.1158/1535-7163.MCT-16-0169-T (2017). Epub 2016 Nov 9.

131. Bocskei, Z., Harmat, V., Vertessy, B. G., Ovadi, J. & Naray-Szabo, G. Calmodulin complexed with trifluoperazine (1:2 complex). RCSB Protein Data Bank, DOI: 10.2210/pdb1A29/pdb (1998). PDB ID: 1A29; deposited 1998-01-19; released 1998-09-16.

132. Vandonselaar, M., Hickie, R. A., Quail, J. W. & Delbaere, L. T. J. Calmodulin complexed with trifluoperazine (1:4 complex). RCSB Protein Data Bank, DOI: 10.2210/pdb1LIN/pdb (1996). PDB ID: 1LIN; deposited 1995-10-11; released 1996-03-08.

133. O’Brien, M. T. et al. Impact of genetic variation on human camkk2 regulation by ca2+-calmodulin and multisite phosphorylation. Sci. Reports 7, 43264, DOI: 10.1038/srep43264 (2017).

134. Ramagopal, U. A., Dulyaninova, N. G., Almo, S. C. & Bresnick, A. R. Structure of s100a4 with pcp. RCSB Protein Data Bank, DOI: 10.2210/pdb3M0W/pdb (2010). PDB ID: 3M0W; deposited 2010-05-12.

135. Broadbelt, K. & Jones, L. B. Evidence of altered calmodulin immunoreactivity in areas 9 and 32 of schizophrenic prefrontal cortex. J. Psychiatr. Res. 42, 612–621, DOI: 10.1016/j.jpsychires.2007.07.006 (2008).

136. Steiner, J., Bielau, H., Bernstein, H. G., Bogerts, B. & Wunderlich, M. T. Increased cerebrospinal fluid and serum levels of s100b in first-onset schizophrenia are not related to a degenerative release of glial fibrillar acidic protein, myelin basic protein and neurone-specific enolase from glia or neurones. J. Neurol. Neurosurg. & Psychiatry 77, 1284–1287, DOI: 10.1136/jnnp.2006.093427 (2006).

137. Emanuele, E. et al. Serum levels of soluble receptor for advanced glycation endproducts (srage) in patients with different psychiatric disorders. Neurosci. Lett. 487, 99–102, DOI: 10.1016/j.neulet.2010.10.003 (2011).

138. Takeda, M. et al. Altered serum glyceraldehyde-derived advanced glycation end product (age) and soluble age receptor levels indicate carbonyl stress in patients with schizophrenia. Neurosci. Lett. 593, 51–55, DOI: 10.1016/j.neulet.2015.03.002 (2015).

139. Nikolac Perkovic, M. et al. Monoamine oxidase and agitation in psychiatric patients. Prog. Neuro-Psychopharmacology Biol. Psychiatry 69, 131–146, DOI: 10.1016/j.pnpbp.2016.02.002 (2016). Epub 2016 Feb 4.

140. Turner, W. J. & Merlis, S. A clinical trial of pargyline and dopa in psychotic subjects. Dis. Nerv. Syst. 25, 538–541 (1964).

141. Saunders, J. C. Treatment of hospitalized depressed and schizophrenic patients with monoamine oxidase inhibitors: including reflections on pargyline. Annals New York Acad. Sci. 107, 1081–1089, DOI: 10.1111/j.1749-6632.1963.tb13351.x (1963).

142. Substance Abuse and Mental Health Services Administration. Impact of the DSM-IV to DSM-5 Changes on the National Survey on Drug Use and Health. [Internet] (2016). Table 3.22, DSM-IV to DSM-5 Schizophrenia Comparison.

143. Kesby, J., Eyles, D., McGrath, J. et al. Dopamine, psychosis and schizophrenia: the widening gap between basic and clinical neuroscience. Transl. Psychiatry 8, 30, DOI: 10.1038/s41398-017-0071-9 (2018).

144. DrugBank. Pizotifen. https://go.drugbank.com/drugs/DB06153 (Accessed 2025). DrugBank ID: DB06153.

145. DrugBank. Loxapine. https://go.drugbank.com/drugs/DB00408 (Accessed 2025). DrugBank ID: DB00408.

146. DrugBank. Clozapine. https://go.drugbank.com/drugs/DB00363 (Accessed 2025). DrugBank ID: DB00363.

147. DrugBank. Olanzapine. https://go.drugbank.com/drugs/DB00334 (Accessed 2025). DrugBank ID: DB00334.

148. Ota, K. Y., Kurland, A. A. & Ferro-Diaz, P. Sch-12041 in the treatment of acute schizophrenic patients. Curr. Ther. Res. Clin. Exp. 15, 327–332 (1973).

149. Fann, W. E., Pitts, W. M. & Wheless, J. C. Pharmacology, efficacy, and adverse effects of halazepam, a new benzodiazepine. Pharmacotherapy 2, 72–79, DOI: 10.1002/j.1875-9114.1982.tb03177.x (1982).

150. Szarmach, J., Włodarczyk, A., Cubała, W. J. & Wiglusz, M. S. Benzodiazepines as adjunctive therapy in treatment refractory symptoms of schizophrenia. Psychiatr. Danub. 29, 349–352 (2017).

151. Kahn, J. P., Puertollano, M. A., Schane, M. D. & Klein, D. F. Adjunctive alprazolam for schizophrenia with panic anxiety: clinical observation and pathogenetic implications. The Am. J. Psychiatry 145, 742–744, DOI: 10.1176/ajp.145.6.742 (1988).

152. DrugBank. Alprazolam. https://go.drugbank.com/drugs/DB00404 (Accessed 2025). DrugBank ID: DB00404.

153. DrugBank. Halazepam. https://go.drugbank.com/drugs/DB00801 (Accessed 2025). DrugBank ID: DB00801.

154. DrugBank. Diazepam. https://go.drugbank.com/drugs/DB00829 (Accessed 2025). DrugBank ID: DB00829.

155. National Center for Advancing Translational Sciences (NCATS). Butaperazine maleate. Inxight Drugs, identifier 22VUW43J2H (1967). Antipsychotic phenothiazine; originator: Bayer; first approved in 1967; previously marketed in the US.

156. Evaluation of a new antipsychotic agent: Butaperazine maleate (repoise maleate). JAMA 206, 2307–2308, DOI: 10.1001/jama.1968.03150100057014 (1968).

